# Brain-controlled augmented hearing for spatially moving conversations in multi-talker environments

**DOI:** 10.1101/2024.02.05.579018

**Authors:** Vishal Choudhari, Cong Han, Stephan Bickel, Ashesh D. Mehta, Catherine Schevon, Guy M. McKhann, Nima Mesgarani

**Affiliations:** Department of Electrical Engineering, Columbia University, New York City, New York, USA; Mortimer B. Zuckerman Mind Brain Behavior Institute, New York City, New York, USA; Hofstra Northwell School of Medicine, Uniondale, New York, USA; The Feinstein Institutes for Medical Research, Manhasset, New York, USA; Department of Neurology, Columbia University, New York City, New York, USA

## Abstract

Focusing on a specific conversation amidst multiple interfering talkers presents a significant challenge, especially for the hearing-impaired. Brain-controlled assistive hearing devices aim to alleviate this problem by separating complex auditory scenes into distinct speech streams and enhancing the attended speech based on the listener’s neural signals using auditory attention decoding (AAD). Departing from conventional AAD studies that relied on oversimplified scenarios with stationary talkers, we present a realistic AAD task that mirrors the dynamic nature of acoustic settings. This task involves focusing on one of two concurrent conversations, with multiple talkers taking turns and moving continuously in space with background noise. Invasive electroencephalography (iEEG) data were collected from three neurosurgical patients as they focused on one of the two moving conversations. We propose an enhanced brain-controlled assistive hearing system that combines AAD and a binaural speaker-independent speech separation model. The separation model unmixes talkers while preserving their spatial location and provides talker trajectories to the neural decoder to improve auditory attention decoding accuracy. Our subjective and objective evaluations show that the proposed system enhances speech intelligibility and facilitates conversation tracking while maintaining spatial cues and voice quality in challenging acoustic environments. This research demonstrates the potential of our approach in real-world scenarios and marks a significant step towards developing assistive hearing technologies that adapt to the intricate dynamics of everyday auditory experiences.

**TAKEAWAYS:**

- Brain-controlled hearing device for scenarios with moving conversations in multi-talker settings, closely mimicking real-world listening environments
- Developed a binaural speech separation model that separates speech of moving talkers while retaining their spatial locations, enhancing auditory perception and auditory attention decoding
- Proposed system enhances speech intelligibility and reduces listening effort in realistic acoustic scenes

## INTRODUCTION

Speech communication in multi-talker environments is challenging, particularly for the hearing impaired^1^. Modern hearing aids, though proficient at suppressing general background noises^2,3^, fall short in a critical aspect: they cannot selectively enhance the attended talker’s speech without first knowing which talker is the target^4^. This limitation underscores the need for a brain-controlled approach, in which the listener’s neural responses are used to decode and enhance the talker to whom attention is directed^5^, a technique known as auditory attention decoding (AAD)^6^. In parallel, the field of automatic speech separation has seen significant progress in the recent years^7–9^. Speech separation aims to isolate individual talkers from a mixture (captured by one or more microphones). Auditory attention decoding can be combined with automatic speech separation to enable a brain-controlled hearing device^6,10–13^ by isolating and amplifying the speech of the attended talker, therefore mitigating the challenges posed by multi-talker environments.

Past studies have established the feasibility of decoding auditory attention from both invasive^5,6,10,11^ and non-invasive^13–15^ neural recordings. Despite these advancements, existing studies predominantly employ overly simplistic acoustic scenes that do not mimic the real world scenarios^6,10–14,16^. Common experimental setups have been limited to stationary talkers without background noise, and primarily focus on distinguishing between two concurrent talkers. This lack of realism in experimental design is a significant barrier to the generalization of these technologies to everyday life scenarios. Real-world listening involves dynamic conversation involving multiple talkers, often engaged in turn-taking while moving in space, all amidst varying background noises. Our study aims to bridge this gap by simulating a more realistic experimental paradigm, therefore advancing the field of AAD towards practical applications. Another important factor that past research has often overlooked is the listeners’ desire to track moving talkers in space. This aspect is crucial for natural listening^17^ and, thus, for the effectiveness of brain-controlled hearing devices. A successful brain-controlled hearing device must separate speech streams as they move in space while preserving the perceived spatial location of each talker^18^. Previous studies of AAD have been based on decoding only the spectro-temporal features of speech. However, recent scientific studies have shown that the human auditory cortex also encodes the location of the attended talker^19,20^ which can potentially lead to the ability to decode the spatial trajectory of attended talkers. Our study takes a crucial step by investigating whether adding talker trajectories can improve the AAD performance.

Another persistent challenge in AAD model fitting and evaluation is the difficulty in accurately determining the specific talker to which a subject is attending, especially with high temporal resolution. Previous methods often assume that subjects continuously focus on a pre-designated talker, overlooking the possibility of inadvertent attention shifts^21^. This assumption can lead to mislabeling in data and biasing the performance evaluation of AAD algorithms. Our study addresses this issue by integrating a behavior measure into our experimental design to ascertain the ongoing focus of the subject more precisely, thereby enhancing the reliability of our data and the validity of our evaluation metrics.

In this work, we present a comprehensive and novel approach to AAD that uses complex, dynamic stimuli that more closely resemble real-world acoustic environments. Specifically, we use two concurrent conversations that feature moving talkers and natural background noise, alongside speaker turn-taking among attended and unattended conversations. Furthermore, we introduce a novel task for determining the ground truth labels in attention-focused conversation by requiring the subject to detect deliberately placed repeated words (1-back task)^22,23^. Lastly, we propose a refined brain-controlled hearing system equipped with a real-time, speaker-independent binaural speech separation model^6,10^ that preserves the spatial location of the talkers and outputs real-time speaker trajectories that are used to increase the decoding accuracy of the attended talker. We demonstrate that the system improves speech intelligibility and conversation tracking and preserves the spatial characteristics and voice quality essential for realistic and immersive auditory experiences. This represents a significant advancement towards brain-controlled hearing devices in real-world listening environments.

## RESULTS

### Subjects and Neural Data

Neural responses from three patients undergoing epilepsy treatment were collected as they performed the task with intracranial electroencephalography (iEEG). Two patients (Subjects 1 and 2) had sEEG depth as well as subdural electrocorticography (ECoG) grid electrodes implanted over the left hemispheres of their brains. The other patient (Subject 3) only had stereo-electroencephalography (sEEG) depth electrodes implanted over their left-brain hemisphere. All subjects had electrode coverage over their left temporal lobe, spanning the auditory cortex. The neural data was processed to extract the envelope of the high gamma band (70-150 Hz) band and was used for the rest of the analysis. Speech-responsive electrodes were determined using t-tests on neural data samples collected during speech v/s silence (see Methods). S1, S2 and S3 had 17, 34 and 42 speech-responsive electrodes respectively as shown in Figure S1.

### Experiment Paradigm

The experiment had a total of 28 multi-talker trials, with the average duration of 44.2 s (standard deviation = 2.0 s) each. As shown in Figure 1a, the trials consisted of two concurrent and independent conversations that were spatially separated and continuously moving in the frontal half of the horizontal plane of the subject. The distances of these conversations from the subject were equal and constant throughout the experiment. Both conversations were of equal power (RMS). Talkers were all native American English speakers. Diotic background noise^24,25^ (either “pedestrian” or “speech babble”) was also mixed along with the conversations at power either 9 dB or 12 dB below the power of a conversation stream.

**Figure 1:**
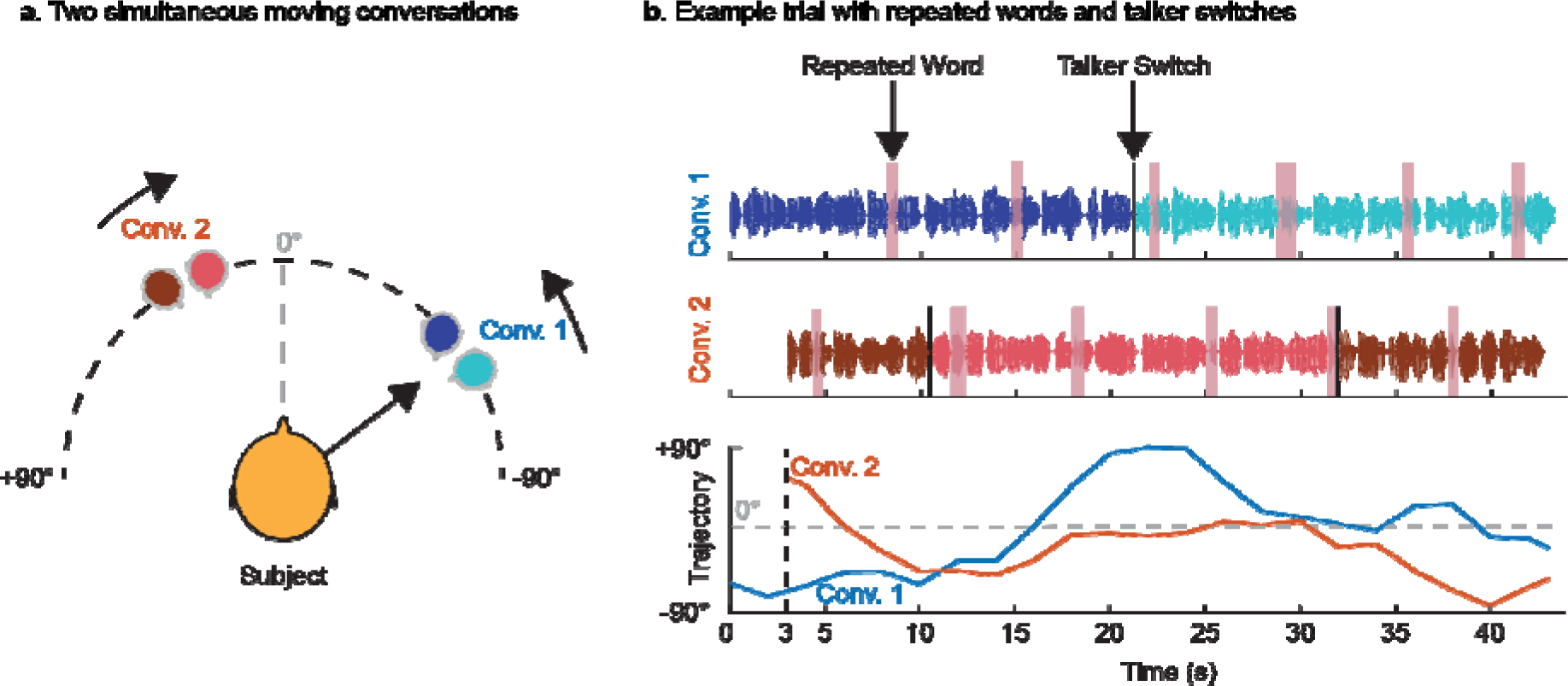
Experiment design. (a) Every trial consisted of two concurrent conversations moving independently in the front hemifield of the subject. Each conversation had two distinct talkers taking turns. (b) Repeated words were inserted across the two conversations as highlighted in pink. The cued (to-be-attended) conversation had a talker switch at 50% trial time mark whereas the uncued (to-be-unattended) conversation had two talker switches, at 25% and 75% trial time marks.

Different talkers took turns in these conversations. As shown in Figure 1b, in the to-be-attended conversation, a talker switch took place at around 50% trial time mark whereas for the to-be-unattended conversation, two talker switches took place, one at around 25% trial time mark and the other nearly at the 75% trial time mark. Repeated words were deliberately inserted in both the conversation streams. The average time interval between the onsets of two repeated words in a conversation was 7.0 s (standard deviation = 1.0 s). The assignment of male and female talkers to various segments of the conversations was counterbalanced across trials to ensure equal durations of concurrent conversations with the same and different genders.

The subjects were instructed to follow (attend to) the conversation that started first and press a push button upon hearing a repeated word in the followed conversation. The uncued (to-be-unattended) conversation started 3 seconds after the onset of the cued (to-be-attended) conversation. The trials were spatialized using head-related transfer functions (HRTFs) and delivered to the subjects via earphones.

### System Proposal

Brain-controlled hearing devices need to combine a speech separation model along with auditory attention decoding to determine and enhance the attended talker. Performing AAD requires having access to individual speech streams and trajectories of every talker in the acoustic scene with which neural representations can be compared to determine the attended talker. Our proposed framework for a binaural brain-controlled hearing device assumes that there are two single-channel microphones, one on the left ear and the other on the right, as shown in Figure 2a. These microphones capture the left and right components of the sounds arriving at the ears of the wearer. The system framework, shown in Figure 2b, makes use of a deep learning-based speaker-independent binaural speech separation model that separates a binaural mixture of speech streams of two moving talkers (recorded by the binaural microphones) into their individual speech streams while also preserving their spatial cues. As spatial cues are preserved in the separated speech streams of the talkers, the model is also able to estimate the trajectories of the moving talkers in the acoustic scene. Auditory attention decoding is enabled by performing canonical correlation analysis which uses the wearer’s neural data and the talkers’ separated speech and estimated trajectory streams to determine and enhance the attended talker by suppressing the unattended talker.

**Figure 2:**
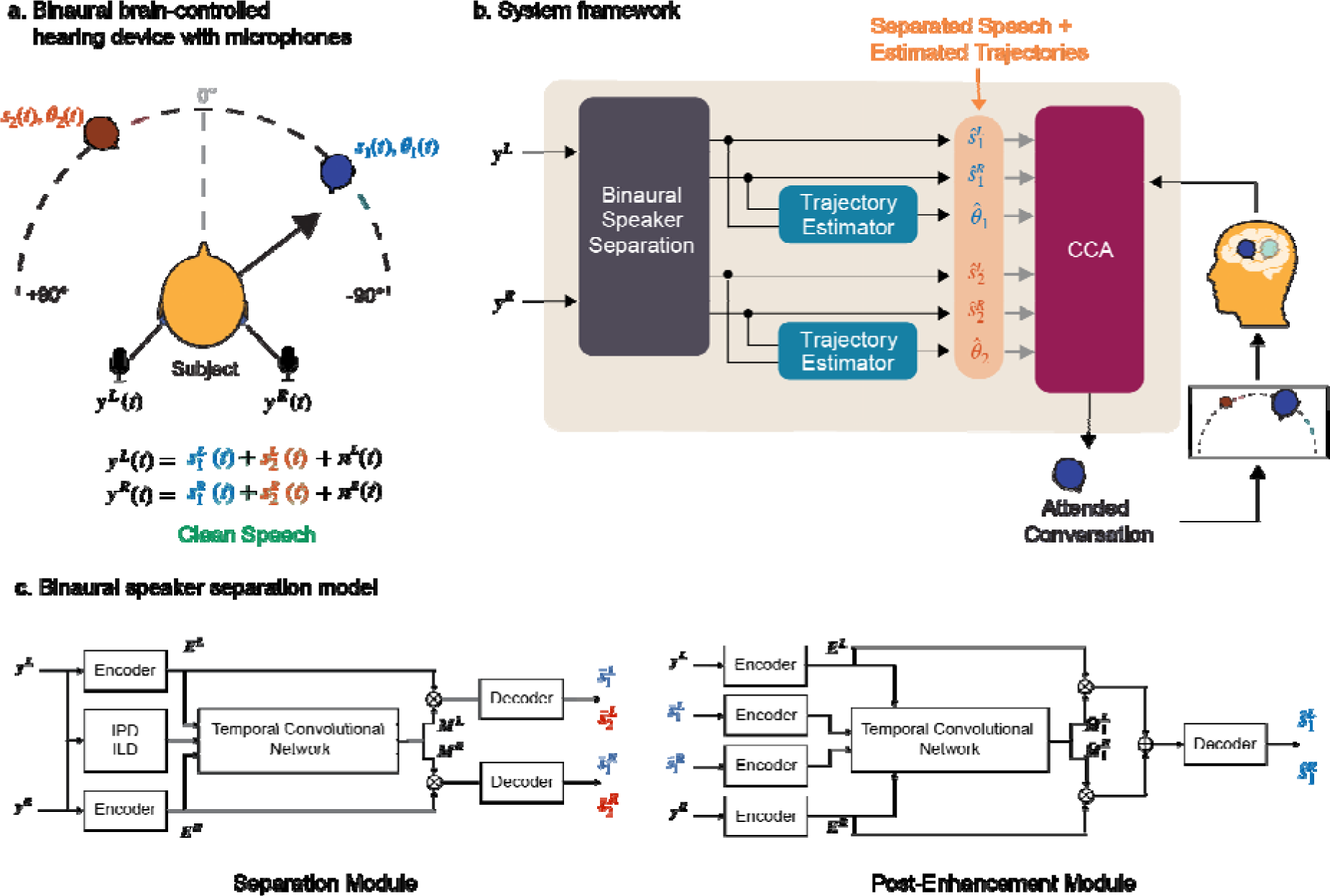
Proposed framework for a binaural brain-controlled hearing device. (a) The framework requires two microphones, one each on both the left and the right ear. The microphones separately capture the left and the right mixtures of sound sources arriving at the ears. (b) The speaker separation works with these microphone recordings to binaurally separate the speech streams while also estimating the trajectories of the talkers. These outputs are used in combination with the wearer’s neural data to decode and enhance the attended talker. (c) The binaural speaker separation model consists of an initial separation module whose outputs are further improved by a post-enhancement module.

### Speaker-Independent Binaural Speech Separation

In this section, we present an automatic speaker-independent speech separation model that separates moving sound sources and preserves spatial cues for all directional sources, enabling listeners to accurately locate each of the moving sources in space. The proposed model, as shown in Figure 2c comprises of two main modules, namely the binaural separation module and the binaural post-enhancement module. Both modules adopt TasNet which has demonstrated exceptional performance in separating audio sources^8,26,27^. Furthermore, the causal configuration enables low-latency processing, making it well-suited for real-time applications.

The binaural separation module takes binaural mixed signals as input and simultaneously separates speech for both left and right channels. Specifically, two linear encoders transform the two channels of mixed signals y^L^, y^R^ ∊ ℝ^T^ into 2-D representations E^L^, E^R^ ∊ ℝ^N×T^, respectively, where T represents the waveform length, N represents the number of encoder bases, and H represents the number of time frames. To explicitly exploit spatial information, we concatenated the encoder outputs and inter-channel phase differences (IPDs) and inter-channel level differences (ILDs) between y^L^ and y^R^, forming spectro-temporal and spatial-temporal features (as detailed in the Materials and Methods section). We then pass this feature through a series of temporal convolutional network (TCN) blocks to estimate multiplicative masks, M^L^, M^R^ ∊ ℝ^CxNxH^ , where C is the number of speakers. In this study, C is prefixed to be two. These multiplicative masks are then applied to E^L^ and E^R^, respectively, and a linear decoder transforms the masked representations back to the waveform of individual speaker, 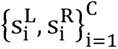. The speaker-independent binaural speech separation module was trained using permutation invariant training^28^. Additionally, we imposed the constraint that the speaker order is the same for both channels, allowing the left- and right-channel signals of each individual speaker to be paired directly. The average signal-to-noise ratio (SNR) improvement of the separated speech over the raw mixture was 14.05 ± 4.79 dB.

The binaural post-enhancement module aims to enhance performance in noisy and reverberant environments because post processing stages have shown effectiveness in improving the signal quality^29^. The module takes each pair of the separated stereo sounds (e. g. , 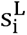 and 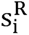) and the mixed signals (y^L^ and y^R^) as input. Similarly, all the encoder outputs are passed through the TCN blocks to estimate multiplicative masks for separating sources. Unlike the speech separation module that only applies multiplicative masks, which is equivalent to spectral filtering, the speech enhancement module performs both multiplication and summation, equivalent to both spectral and spatial filtering. This is similar to multichannel Wiener filtering^30^. Because the input stereo sound (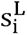 and 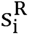) contains both spectral and spatial information of the speaker i, the enhancement module essentially performs informed speaker extraction without the need for permutation invariant training. The average SNR improvement of the enhanced speech over the raw mixture was 16.77 ± 4.92 dB.

The key ingredient of the training is using the signal-to-noise ratio (SNR) as the objective function for both the speech separation and enhancement modules. 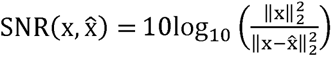, where x and x ^ are the ground truth and estimated waveform, respectively.

Because SNR is sensitive to both time shift and power scale of the estimated waveform, it can force the interaural level difference (ILD) and interaural phase difference (IPD) to be preserved in the estimated waveform. Moreover, we performed utterance-level training on a moving speaker dataset, which encourages the model to leverage spectral and spatial features of speakers in a large context and forces the model to track speakers within the utterance without the need for explicit tracking modules^18^. This approach enables the model to handle moving sources effectively.

We also trained a speaker localizer module using a similar architecture to the enhancement module. The module performs classification of the direction of arrival (DOA) every 80 milliseconds. So, the localizer module estimates a moving trajectory for each moving source which can be utilized to improve the accuracy of attentional decoding. The average DOA error of the estimated trajectories was 4.20 ± 5.76 degrees.

### Behavioral Data Analysis

The push button responses of subjects to repeated words in the conversation being followed help in determining to which conversation a subject was attending. A repeated word in a conversation was considered as correctly detected only if a button press was captured within two seconds of its onset. As shown in Figure S2, all subjects tracked more than 65% of the repeated words in the cued (to-be-attended) conversation. We assign these as hits. However, we see that subjects also tracked a non-zero fraction of repeated words in the uncued (to-be-unattended) conversation (false alarms) indicating that there might have been occasions when the subjects were attending to the uncued (to-be-unattended) conversation. We combined the hit rate and false alarm rate for each subject to generate a sensitivity index (d’) inspired by signal detection theory^22,31^ (SDT). Sensitivity index for each subject was calculated as: d’ = z(False Alarm Rate) – z(Hit Rate), where z(x) is the z-score corresponding to the right-tail p-value of x^31^. Subjects were ranked based on their sensitivity indices (S1: 2.8, S2: 2.3, S3: 1.9).

### Auditory Attention Decoding

In order to decode the attended talker, neural signals were compared with speech spectrograms and trajectories of talkers using canonical correlation analysis^32^ (CCA) (see Methods). For certain trials where it was evident that the subject was following the uncued conversation (greater than or equal to two repeated words detected in the uncued stream), the “attend” and “unattend” labels were swapped for the conversations in that portion. Subject-wise CCA models were trained, and their performance was evaluated using leave-one-trial-out cross validation, i.e., training on N -1 trials and testing on the windows from the N^th^ trial. During training, the CCA models simultaneously learn forward filters on attended talker’s clean speech spectrogram and trajectory and backward filters on the neural data such that upon projection with these filters, the neural data and the attended talker stimuli would be maximally correlated. During testing, these learnt filters are applied to the neural data as well as to every talker’s speech spectrogram and trajectory. The talker which yields the highest correlation score (based on voting of the top three canonical correlations) was determined as the attended talker. We chose a receptive field of 500 ms for neural data and 200 ms for stimuli spectrograms and trajectories (see Methods). The starting sample of the receptive field windows were aligned in time for both neural data and stimuli.

We evaluated auditory attention decoding accuracies for all subjects for a range of window sizes from 0.5 s to 32 s for the following two stimuli versions:

1. Clean Stimuli: Using the clean (before mixing) ground truth speech spectrograms and trajectories of individual talkers in the acoustic scene.
2. Automatically Separated Stimuli: Using the speech spectrograms and estimated trajectories of talkers yielded by the binaural speech separation model.

Figure 3a shows the attended talker decoding accuracies averaged across subjects as a function of window size for both clean and separated versions after correcting for behavior. For both versions, the attended talker decoding accuracies increase as a function of window size. This is expected since with larger window sizes, more information is available to determine the attended talker. Stimuli version had a very small effect on the AAD accuracies across subjects and window sizes (two-sided Wilcoxon signed-rank test, z = 1.50, p-val = 0.13). This indicates that the AAD performance with automatically separated stimuli is as good as the performance with original clean stimuli (Figure 3a), confirming the efficacy of the proposed speaker-independent speech separation module.

**Figure 3:**
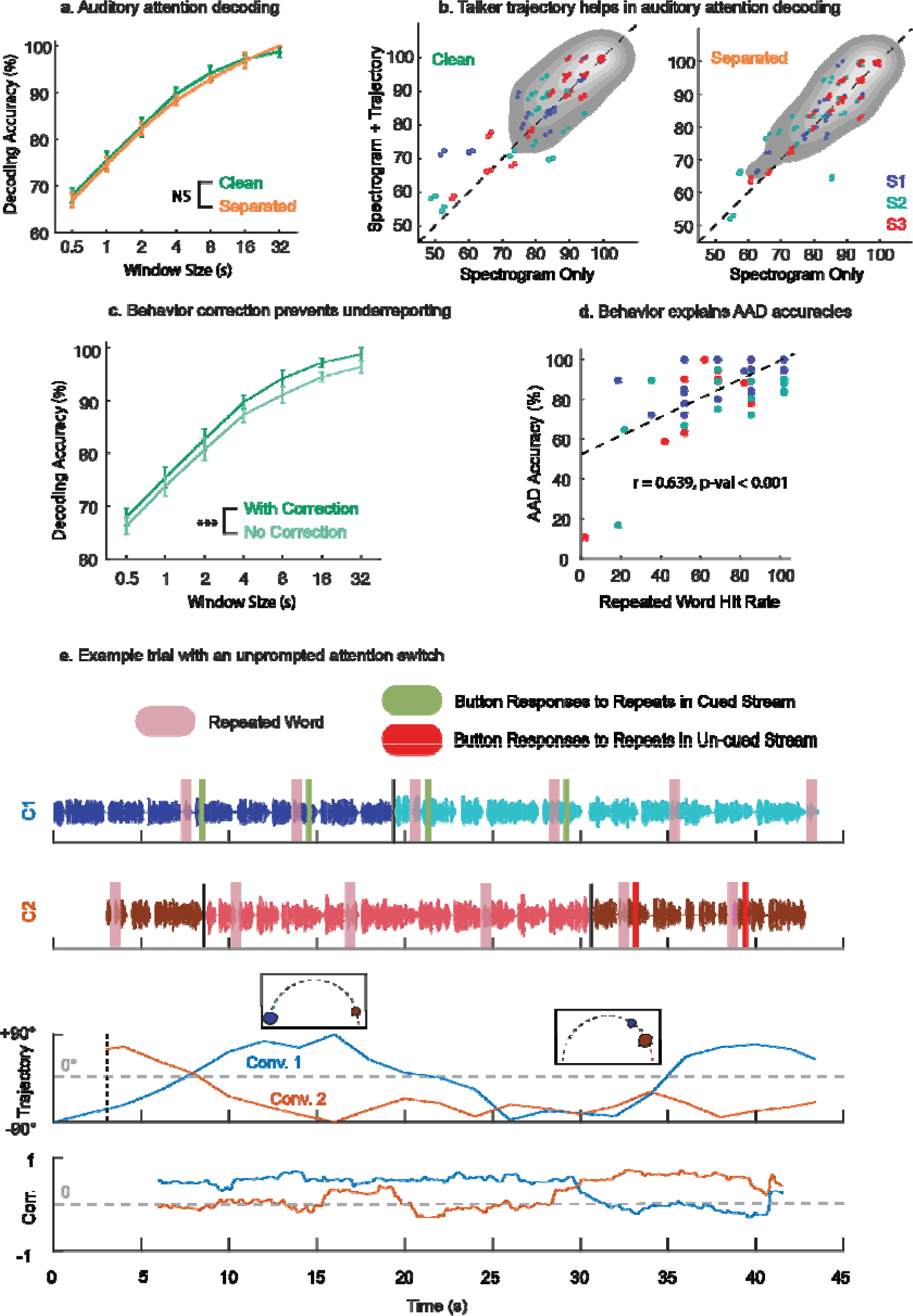
Evaluating auditory attention decoding (AAD) performance and its correlation with behavior. (a) AAD accuracies averaged across subjects as a function of window size. The decoding accuracies are comparable between the clean and separated versions (two-sided Wilcoxon signed-rank test, z = 1.50, p-val = 0.13). Error bars indicate the standard error of mean. (b) Scatter plots comparing trial-wise AAD accuracies for a window size of 4 s when using only spectrogram vs spectrogram + trajectory. Each point represents a trial. AAD accuracies improved significantly when talker trajectories were also incorporated in addition to their speech spectrograms for both clean (two-sided paired t-test, t = 3.2235, df = 80, p-val = 0.002, 95% CI: 0.7534 to 3.1845) and separated (two-sided paired t-test, t = 2.6316, df = 80, p-val = 0.010, 95% CI: 0.3470 to 2.4995) versions. (c) A separate set of models were trained without correcting for behavior. The decoding accuracies are plotted for the clean version of speech for both with and without behavior correction. Not correcting for behavior can lead to significant underreporting of AAD performance (two-sided Wilcoxon signed-rank test, signed-rank = 0, p-val < 0.001) (d) For models trained without correcting for behavior, trial-wise behavioral performance and AAD accuracies are significantly correlated (Pearson’s r = 0.639, p-val < 0.001). (e) An example trial from one of the subjects who shifts attention from the cued conversation (Conv. 1) to the uncued conversation (Conv. 2) in the middle of the trial. Repeated words in the conversation streams are shaded in pink. Button press responses to the repeated words are shown in green (red) for the cued (uncued) conversation. The last plot shows the first canonical correlation for both the conversation streams obtained by continuously sliding a 4 s window. Behavior is well correlated with the canonical correlations.

We studied the improvement in AAD performance when talker trajectories are included in addition to talker spectrograms. For this comparison, we trained and tested CCA models (post behavior correction) with only talker spectrograms without trajectories. As shown in Figure 3b, we found that trial-wise AAD performance improved when talker trajectories were also incorporated in addition to talker spectrograms for both clean (two-sided paired t-test, t = 3.2235, df = 80, p-val = 0.002, 95% CI: 0.7534 to 3.1845) and automatically separated (two-sided paired t-test, t = 2.6316, df = 80, p-val = 0.010, 95% CI: 0.3470 to 2.4995) versions of the stimuli.

Lack of having a behavioral measure and not correcting for the same can lead to underreporting of AAD performance. To study this, we also trained a set of CCA models assuming that the subjects always paid attention to the cued (to-be-attended) conversation. Figure 3c compares the AAD performance for clean stimuli when correcting and not correcting for behavior. Not correcting for behavior significantly hurts AAD performance (two-sided Wilcoxon signed-rank test, signed-rank = 0, p-val < 0.001). This is also true when evaluating with the automatically separated version of the stimuli (two-sided Wilcoxon signed-rank test, signed-rank = 0, p-val < 0.001).

Next, for the models trained without correcting for behavior, we examined whether the behavioral performance on the repeated word detection task could explain the AAD performance on a trial-by-trial basis. We first computed the proportion of repeated words detected in the cued conversation (hit rate) for each trial and for each subject. We also computed corresponding trial-wise AAD accuracies for a window size of 4 s. As shown in Figure 3d, we found that hit rate on the repeated word detection task was significantly correlated with the trial-wise AAD accuracies (Pearson’s r = 0.639, p-val < 0.001). Figure 3e shows an example trial from one of the subjects who, based on behavioral responses, was initially attending to the cued (to-be-attended) conversation and then later attends to the uncued (to-be-unattended) conversation after the conversations cross in space. The canonical correlations mapping the neural data with both the cued and uncued stimuli also capture this shift of attention from one conversation to the other. Thus, the repeated word detection task helps explain AAD performance on a trial-by-trial basis.

### System Dynamics During Talker Transitions

Turn-takings during conversations create talker switches in the attended conversation. For good user experience, it is important that the system tracks the talker switch and seamlessly enhances the new attended talker. Our experiment paradigm, inspired by real-world settings, had asynchronous talker switches in both to-be-attended and to-be-unattended conversations. The new talker continued at the same location as the previous talker in the conversation. The speaker separation model was able to put talkers of a conversation on the same output channel using location and talker continuity. As a result, the system was able to seamlessly track turn-takings in conversations, as shown in Figure 4a.

**Figure 4:**
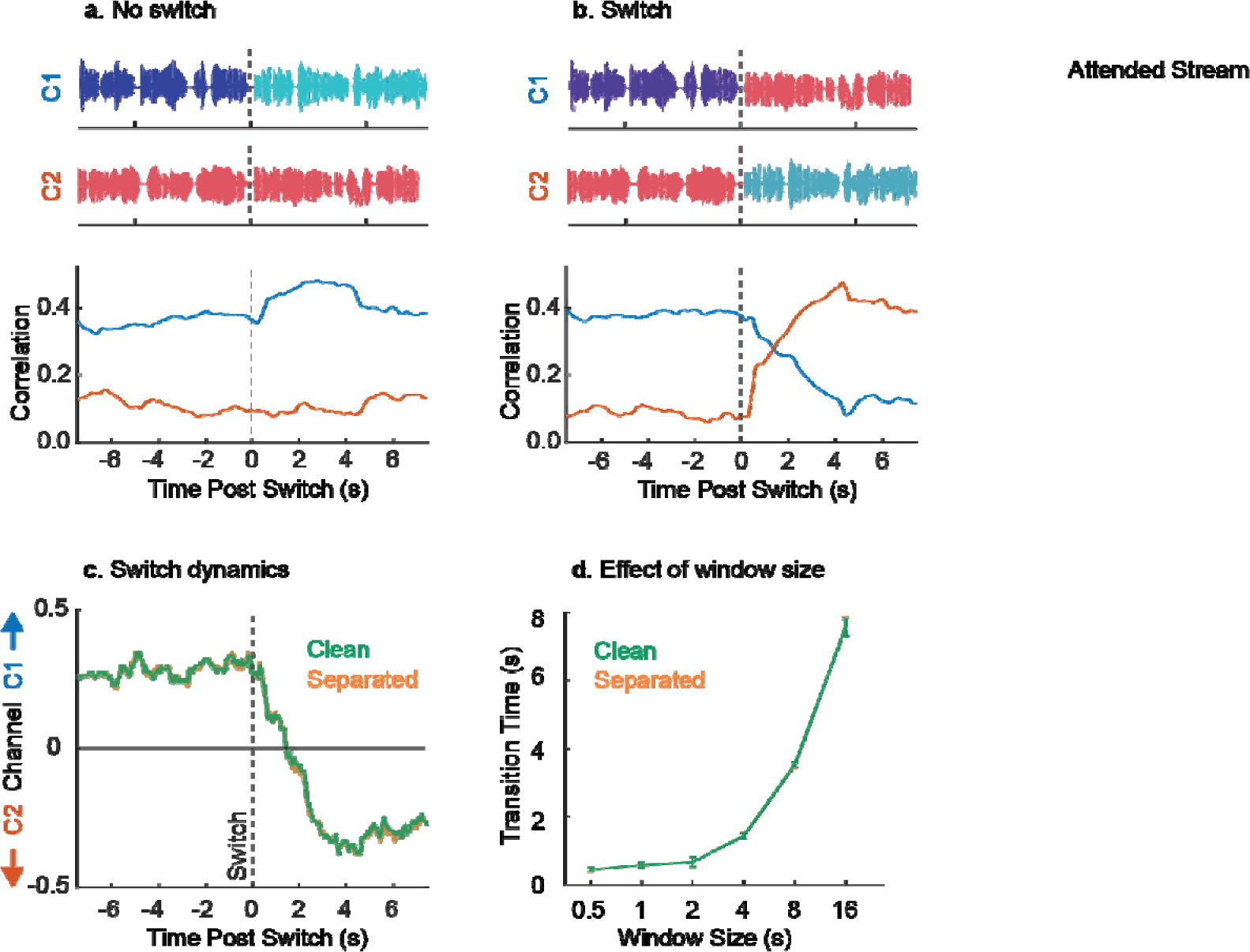
(a) The proposed system seamlessly tracks turn-takings. This is facilitated by the speaker separation module which places talkers in a conversation on the same output channel by relying on location and talker continuity cues. Attended conversation is highlighted with a pink shade. Correlations showed are the average of the top three canonical correlations for separated version of the stimuli. (b) Attention switch from one conversation to another can be simulated by swapping the output channels of the binaural separation system. (c) Channel preference dynamics after simulated attention switch for a decoding window size of 4 s. (d) Transition times as a function of decoding window size. No significant differences were observed between the clean and separated versions (two-sided Wilcoxon signed-rank test, signed-rank = 17, p-val = 0.70). Error bars in all plots indicate the standard error of mean.

In some cases, the wearer of the hearing device might switch attention from a conversation at a particular location to another conversation at a different location. To study how our system responds in such cases, we artificially swapped the outputs of the binaural speech separation system at the point of talker switch in the cued conversation, as shown in Figure 4b. Since we combine the results of the top three canonical correlations based on voting to determine the attended talker or channel, we define a metric channel preference index (CPI), i.e.,

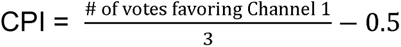

Thus, a positive CPI would indicate a preference to Channel 1 whereas a negative CPI would indicate a preference to Channel 2. In Figure 4c, we show the CPI averaged across trials for one of the subjects (S3) when attention switch is simulated. We define the transition time as the time point where the average CPI crosses 0. Figure 4d shows the transition times (averaged across subjects) as a function of window size for both clean and separated versions. No significant difference was found in the transition times across subjects and window sizes between the clean and separated versions (two-sided Wilcoxon signed-rank test, signed-rank = 17, p-val = 0.70).

### Evaluation of System Performance

#### >Part A: Subjective

To evaluate the performance of the proposed system, an online Amazon MTurk experiment was conducted with 24 native speakers of American English with self-reported normal hearing. The participants listened to simulated output of the proposed system using the neural signatures obtained by concatenating the channels from all the three subjects, for a total of 15 trials, five for each of the following conditions, in a blind fashion:

1. System Off: The raw mixture stimuli that was played to the subjects from whom neural data was recorded.
2. System On (Separated): Mixture in which the attended talker, as determined by the neural signatures, was enhanced using the output of the binaural speaker-separation model.
3. System On (Clean): Mixture in which the attended talker was enhanced using clean ground truth speech.

Enhanced mixtures were generated by suppressing the un-attended talker and the background noise in the mixture by the same scale factor such that the resulting power difference between the attended and the unattended talker was 9 dB. Like the iEEG participants, the MTurk participants were also instructed to follow the cued conversation and press space bar on their keyboards upon hearing the repeated words in the conversation being followed. After each trial, the participants were asked to rate the difficulty of following the cued conversation on a scale from 1 (very difficult) to 5 (very easy) and the quality of voices in the conversation on a scale from 1 (bad) to 5 (excellent). To also test the intelligibility of the conversations, the participants were also asked to respond to a multiple-choice question based on the content of the cued conversation after each trial. A short localization task was also included at the end of each trial to determine if the attended talker can be localized post enhancement.

Figure 5 summarizes the results. As shown in Figure 5a, under both the “system on” conditions, the repeated word detection accuracy in the cued conversation is enhanced when compared to the “system off” condition (two-sided paired t-test, p-val < 0.001), whereas for the uncued conversation (Figure 5b), the detection accuracy is reduced (two-sided paired t-test, p-val < 0.01). This means that the system helps track the cued conversation and prevents unintentional tracking of the uncued conversation. We also find that intelligibility of the cued conversation is significantly enhanced under the “system on” conditions (Figure 5c, two-sided paired t-test, p-val < 0.05). No significant differences are observed between the clean and separated versions of the “system on” condition. Ease of attending to the cued conversation increases from “system off” condition to “system on with separated speech” condition (two-sided paired t-test, p-val < 0.0001) to “system on with clean speech” condition (two-sided paired t-test, p-val < 0.01), as shown in Figure 5d. Surprisingly, no differences in voice quality of the talkers in the cued conversation were observed between the “system off” and the “system on with separated speech” condition (Figure 5f). However, participants rated the voice quality in the “system off with clean speech” condition higher than the other two (two-sided paired t-test, p-val < 0.05). These results indicate that a scope for improvement exists for the speaker-independent binaural speech separation model and its upper bounds (when there is ideal separation) are captured by the “system on with clean speech” condition. The ability to localize talkers in space, as shown in Figure 5e, was comparable across all the three conditions highlighting retention of the attended talker spatial cues when the system is turned on. In summary, the system helps follow the conversation of interest, increases its intelligibility and the ease of attending to it while also preserving spatial cues.

**Figure 5:**
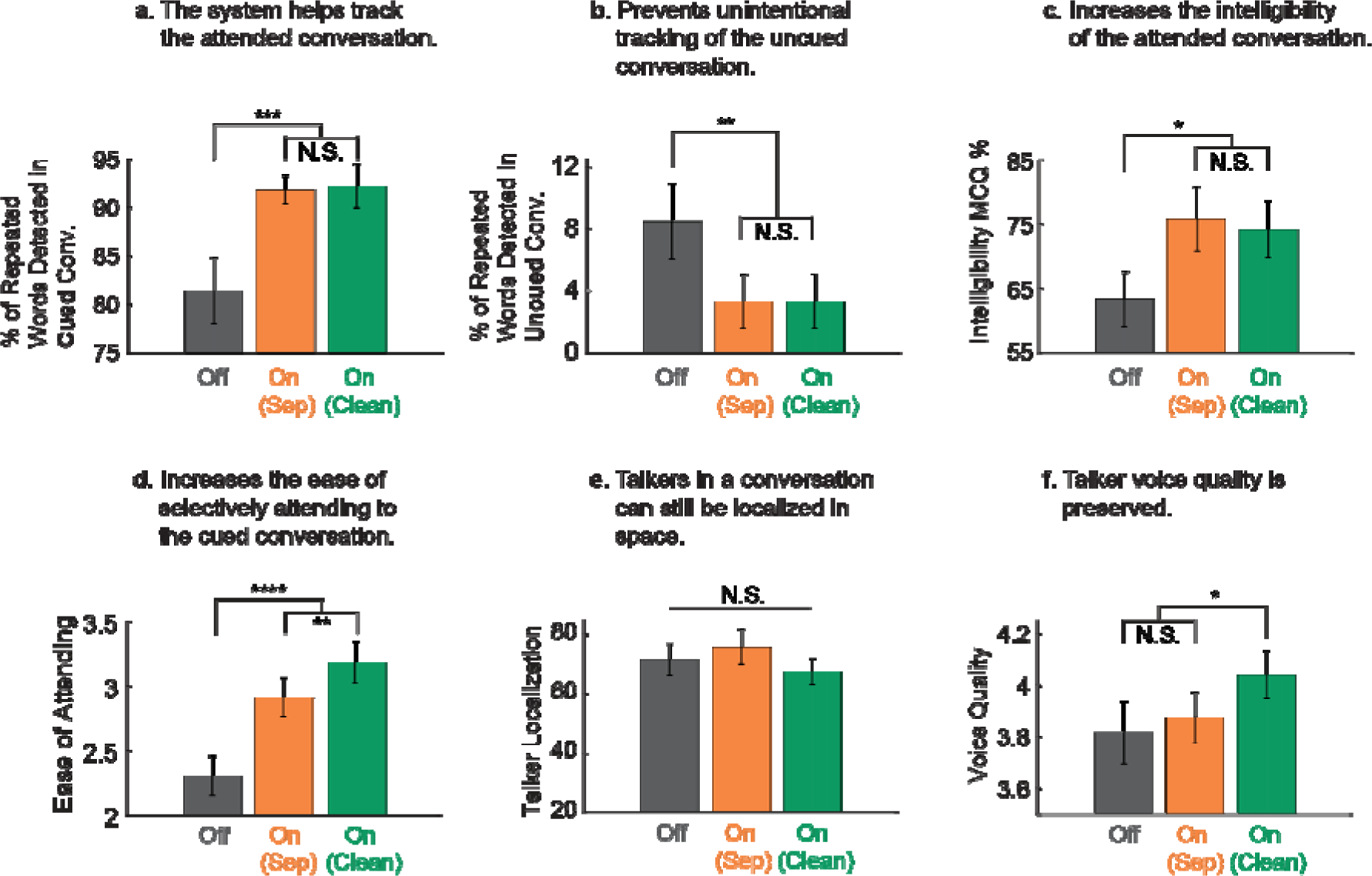
Subjective evaluation of system outputs show enhanced tracking of the cued conversation, improved intelligibility and retention of talker cues and voice quality. Twenty-four online participants listened to trials from the following conditions: (i) System Off: raw original mixture played to iEEG subjects, (ii) System On (Separated): attended talker enhanced with the output of the binaural separation system, (iii) System On (Clean): attended talker enhanced with clean ground truth speech. (a) Repeated word detection accuracy in the cued conversation increases significantly when the system is turned on for both clean as well as separated versions (two-sided paired t-test, p-val < 0.001). (b) Repeated word detection accuracy for the uncued conversation drops significantly when the system is turned on (two-sided paired t-test, p-val < 0.01). (c) Intelligibility of the cued conversation is significantly increased under the system on conditions (two-sided paired t-test, p-val < 0.05) (d) Attending to the cued conversation is easier under the system on conditions (two-sided paired t-test, p-val < 0.0001). (e) Participants can localize talkers in space equally well in all conditions. (f) No significant difference in voice quality ratings was observed between the system off condition vs the system on with separated speech condition. However, participants rated the voice quality of the system on with clean speech condition to be relatively higher (two-sided paired t-test, p-val < 0.05). Error bars in all plots indicate the standard error of mean.

#### >Part B: Objective

In addition to subjective evaluation, we also performed an objective evaluation where the same system simulated outputs in the subjective evaluation were compared with their corresponding clean to-be-attended conversation waveforms (as reference) to calculate narrowband MOS-mapped Perceptual Evaluation of Speech Quality^33^ (PESQ) and Extended Short-Time Objective Intelligibility^34^ (ESTOI) scores. As expected, in Figure 6, we see a significant improvement in these scores as we progress from “system off” condition to “system on with separated speech” condition to “system on with clean speech” condition (two-sided paired t-tests, p-val < 0.0001).

**Figure 6:**
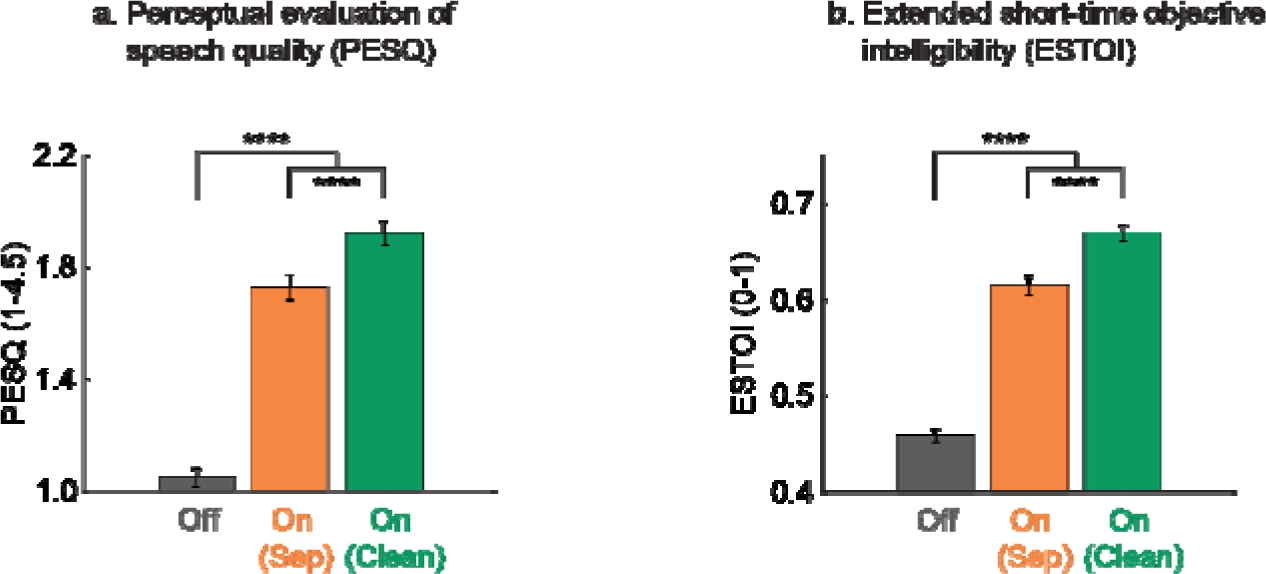
Improved objective quality and intelligibility. (a and b) Both PESQ and ESTOI scores increase from “system off” condition to “system on with separated speech” condition to “system on with clean speech” condition (two-sided paired t-tests, p-val < 0.0001). Error bars in all plots indicate the standard error of mean.

## DISCUSSION

We introduced a novel AAD experimental paradigm that diverges from existing studies by incorporating concurrent conversations with natural turn-takings where talkers move in space amidst background noise. This approach represents a substantial advancement in creating realistic auditory scenarios for AAD research. Our binaural speaker separation system successfully separated these dynamic conversations into individual streams while preserving talker spatial cues. Additionally, the speech separation system provides real-time talker trajectories to the AAD algorithm, enhancing its decoding accuracy. The use of the repeated word detection task across the conversations provided a robust ground truth label for the attended conversation with a high temporal resolution and explained AAD performance on a trial-by-trial basis. Evaluations of the proposed system revealed improved tracking of the attended conversation and increased its intelligibility while preserving the perceived location of each talker in space.

The primary aim of this study was to address the limitations of previous AAD research that predominantly assumed two stationary talkers^6,10,11,13,14^, thereby restricting the applicability of such research to real-world scenarios. In realistic acoustic scenes, we normally listen to simultaneous conversations which can involve multiple talkers. Our research extends previous work by replacing concurrent talkers with concurrent conversations involving natural turn-taking. By introducing a speaker-independent speech separation model that leverages both spatial and spectro-temporal information, our research marks a significant step toward creating an immersive listening experience that closely mimics natural environments. This model not only separates the speech of moving talkers but also allows listeners to accurately track their locations, an aspect crucial for realistic AAD applications. An essential contribution of our study is that incorporating real-time talker trajectories estimated by the speech separation algorithm in addition to spectro-temporal information can improve AAD accuracy^19,35–37^. Further research is needed to distinguish listener motion-induced from talker motion-induced acoustic change and how it could be encoded differently in the human auditory cortex^38,39^.

Another contribution of our study is introducing a behavioral task of repeated word detection across conversations, allowing us to identify the actual attended conversation with high temporal resolution. This method addresses a common issue in previous AAD studies where subjects’ attention could inadvertently shift to the unattended stream^21^, leading to mislabeled data and affecting the training and evaluation of AAD models. By incorporating a behavioral measure into our experiment design, we have enhanced the accuracy of determining the attended talker or conversation. In future AAD studies with moving talkers, a higher degree of temporal resolution can be achieved by asking the subjects also to report the spatial trajectory of the conversation followed. Additionally, further research is needed to investigate the difference between endogenous and exogenous auditory attention switches and how they may be decoded differently^40^.

While our study focused on neural activity in the high gamma band, incorporating low-frequency neural activity, which has been shown to track motion and attention, could improve AAD accuracies. Prior invasive^41^ and non-invasive^13,42^ AAD studies have shown signatures of auditory attention (via tracking of the envelope of the attended speech) in the lower frequencies (1 – 7 Hz). A recent study^43^ also showed that low-frequency neural activity also tracks the location of the attended talker, especially in delta (< 2 Hz) phase and alpha (8-12 Hz) power.

Including low-frequency neural signals might provide a more comprehensive understanding of the neural underpinning of auditory attention and enhance the performance of AAD systems.

A critical aspect of future research should involve transitioning to a real-time, closed-loop system. This requires the integration of speech separation and AAD components to work synchronously in a causal, real-time manner. Furthermore, determining how to optimally manipulate the acoustic scene based on the decoded attended talker remains an area for further investigation. Such acoustic modifications should help the listener follow the attended conversation while still maintaining the ability to switch to the unattended one. Our experiment design could be further aligned with real-world scenarios by introducing more complex motion patterns for talkers, such as radial motion and motion pauses. This would add a layer of complexity to the auditory scene, presenting conversations with time-varying power and potentially challenging the current speaker separation model. Addressing this challenge may involve retraining or fine-tuning the model on datasets with these characteristics.

A brain-controlled hearing device that can quickly and accurately adapt to changes in the listener’s attention is a challenge that may be more effectively addressed with invasive neural recording techniques. However, a critique of our approach is the reliance on invasive neural recordings which might be perceived as less accessible. Considering the rapid advancements in speech BCI research involving invasive neural recordings^44–48^, these methods are becoming increasingly common and feasible. The precision and speed offered by invasive recordings are currently unmatched by non-invasive techniques, making them essential for exploring the upper limits of AAD performance. While future research continues to explore less invasive or alternative neural recording methods, our current focus on invasive recordings is crucial for advancing the field and setting benchmarks for performance of these systems and establishing minimum required performance for listeners to prefer AAD functionality.

Our study contributes significantly to AAD research and brain-controlled hearing devices by introducing more realistic experimental paradigms and advancing the technology toward practical applications. The insights from this research enhance our understanding of auditory attention in complex environments and pave the way for future innovations in assistive hearing technologies.

## METHODS

### Participants

The study had a total of three human participants of which two (Subjects 1 and 2) were from North Shore University Hospital (NSUH) and one (Subject 3) was from Columbia University Irving Medical Center (CUIMC). All participants were undergoing clinical treatment for epilepsy. Subjects 1 and 2 were both implanted with subdural electrocorticography (ECoG) grid and stereo-electroencephalography (sEEG) depth electrodes on their left-brain hemispheres.

Subject 3 only had sEEG depth electrodes implanted over their left-brain hemisphere. The electrode targets for these participants were determined purely based on clinical requirements. The participants provided informed consent as per the local Institutional Review Board (IRB) regulations.

### Neural Data Pre-Processing + Hardware

The neural data of participants from NSUH (Subjects 1 and 2) were recorded using Tucker-Davis Technologies (TDT) hardware using a sampling rate of 1526 Hz. The neural data of the participant from CUIMC (Subject 3) was recorded using Natus Quantum hardware using a sampling rate of 1024 Hz. Left and right channels of the audio stimuli played to the participants were also recorded in sync with neural signals to facilitate segmenting of neural data into trials for further offline analysis.

Neural data was pre-processed and analyzed using MATLAB software (MathWorks). All neural data was first resampled to 1000 Hz and then montaged to a common average reference to reduce recording noise^49^. The neural data was then further downsampled to 400 Hz. Line noise at 60 Hz and its harmonics (up to 180 Hz) were removed using a notch filter. The notch filter was designed using MATLAB’s *fir2* function and applied using *filtfilt* with an order of 1000. In order to extract the envelope of the high gamma band (70 - 150 Hz), the neural data was first filtered with a bank of eight filters, each with a width of 10 Hz, spaced consecutively between 70 Hz and 150 Hz^50^. The envelopes of the outputs of these filters were obtained by computing the absolute value of their Hilbert transform. The final envelope of the high gamma band was obtained by computing the mean of the individual envelopes yielded by the eight filters and further downsampling to 100 Hz.

Speech responsive electrodes were determined by comparing neural samples (of the high gamma envelope sampled at 100 Hz) recorded in response to speech with those recorded in response to silence. For each trial, 25 samples corresponding to silence were randomly chosen in a [t = -0.4 s to -0.1 s] window, with t = 0 s being the onset of speech. Similarly, 25 samples corresponding to speech were drawn from a window [t = 0.1 s to 1.6 s]. These samples were accumulated across trials and a t-statistic (between speech and silence samples) was computed for every electrode. Electrodes with a t-statistic above 5 were considered to be speech-responsive and retained for further analysis.

### Stimuli Design and Experiment

The experiment consisted of 28 multi-talker trials with a mean trial duration of 44.2 s (SD = 2.0 s). The total experiment lasted 26 minutes. Every trial consisted of two concurrent and independent conversations (one to-be-attended, one to-be-ignored) that were spatially separated and continuously moving in the presence of diotic background noise. The to-be-ignored conversation started 3 s later than the to-be-attended conversation. The participants were cued to attend to the conversation that started first.

A total of eight native American English voice actors (four male, four female) were recruited to voice these conversations. These conversations were based on general daily life situations (see Supplementary Table S1 for conversation transcripts). Every trial consisted of four talkers: two for the to-be-attended conversation (say A and B), two for the to-be-unattended conversation (say C and D). The to-be-attended conversation had one turn-taking (talker switch) at the 50% trial time mark whereas the to-be-ignored conversation had two turn-takings: one at the 25% trial time mark and the other at the 75% trial time mark. Thus, the talker in the to-be-attended conversation would transition from A to B and the talker in the to-be-ignored conversation would transition from C to D to back to C.

To check to which conversation a participant might be attending, repeated words were artificially inserted in both the to-be-attended and the to-be-ignored conversations. Participants were asked to press a button upon hearing a repeated word in the conversation that they were following. The conversation transcripts were force aligned with the audio recordings of the voice actors using the Montreal Forced Aligner tool^51^. The repeated words were inserted in the conversations based on the following criteria:

- The number of repeated words to be inserted in a conversation of a trial was determined by dividing the trial duration (in seconds) by 7 and rounding the result.
- For every trial, an equal number of repeated words were inserted in the to-be-attended and the to-be-ignored conversations.
- A word could be repeated only if its duration was at least 300 ms.
- To make repeated words sound smooth and natural, a Hanning window of 30 ms was applied to both sides of the audio segment corresponding to the repeated word.
- The audio segment corresponding to a repeated word was also prefixed and postfixed with 200 ms of silence.
- The time interval between the onsets of two repeated words in a conversation was constrained to lie between 5.5 s to 9.5 s.
- There was always one repeated word whose onset was within 1.5 s post talker switch in the to-be-attended conversation. This was done to check if participants tracked the switch in talkers in the to-be-attended conversation.
- The onset of the first repeated word in a trial was constrained to lie between 5 - 8 s from trial start time. This first repeated word could occur either in the to-be-attended conversation or the to-be-ignored conversation.
- The minimum time gap between a repeated word onset in the to-be-attended conversation and a repeated word onset in the to-be-ignored conversation was set to be at least 2.5 s. This was done to prevent simultaneous overlap of repeated words in the two conversations and to allow for determining to which conversation a participant was attending to.

Google Resonance Audio software development kit (SDK) was used to spatialize the audio streams of the conversations^52^. The trajectories for these conversations were designed based on the following criteria:

- The trajectories were confined to the frontal half of the horizontal plane of the subject in a semi-circular fashion. In other words, the conversations were made to move on a semi-circular path at a fixed distance from the subject spanning -90 degrees (right) to +90 degrees (left).
- The trajectories were initially generated with a resolution of 1 degree and a sampling rate of 0.5 Hz using a first order Markov chain.
- This Markov chain had 181 states (-90 degrees to +90 degrees with a resolution of 1 degree). All states were equally probably of being the initial state.
- The subsequent samples of a trajectory were generated with a probability transition matrix shown in Supplementary Figure S3.
- The resulting trajectories were smoothed with a moving average of five samples and then stretched to span the whole frontal half plane.
- The trajectories were further upsampled using linear interpolation to 10 Hz.
- A pair of trajectories corresponding to a pair of conversations in a trial also followed the following criteria:

○ The spatial separation between the conversations when the second conversation starts was set to be at least 90 degrees.
○ The spatial separation between the conversations during the talker switch in the to-be-attended conversation was ensured to be at least 45 degrees.
○ The correlation of the two trajectories were ensured to be less than 0.5.
- A total of 1000 trajectory sets (each with 28 pairs, one for each of the 28 trials) were generated based on the above criteria.
- To have the trajectories span a uniform joint distribution, the set with the highest joint entropy (computed with a bin size of 20 degrees) was chosen as final.

In addition to the two conversation streams, a single channel background noise was duplicated for both left and right channels introduced in the auditory scene. For every trial, the background noise was either pedestrian noise^24^ or speech babble noise^25^. When mixing the three streams the power of the two conversation streams were always kept the same. The power of the background noise stream was suppressed relative to the power of a conversation stream by either 9 dB or 12 dB. Trial parameters such as background noise type, its power level and voice actor assignments were all counterbalanced across the trials. Stimuli was delivered to the participants with a sampling rate of 44.1 kHz through stereo earphones (Panasonic RP-HJE120).

### Speaker-Independent Binaural Speech Separation

#### Cross-Domain Features

Although the encoder outputs E^L^ and E^R^ contain both spectral and spatial information, we added interaural phase difference (IPD) and interaural level difference (ILD) as additional features to increase speaker distinction when speakers are at different locations^53,54^. Specifically, we calculated cos(IPD), sin(IPD), and ILD ∊ ℝ^F×H^,

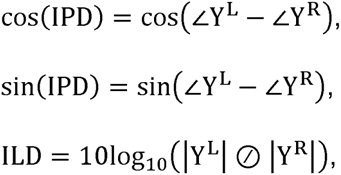

where Y^L^, Y^R^ ∊ ℝ^F×H^ are the STFT output of Y^L^, Y^R^, respectively, F is the number of frequency bins, and ⊘ is element-wise division operation. The hop size for calculating Y^L^ and Y^R^ is the same as that for E^L^ and E^R^ to ensure they have the same number of time frames, even though the window length in the encoder is typically much shorter than that in the STFT. Finally, we concatenated these cross-domain features into[E^L^, E^R^, cosCIPD), sinCIPD), ILDJ ∊ ℝ^(2N+3F)×H^ as the input to the binaural speech separation module.

#### Training and Development Datasets

For the training and development sets, we generated 24,000 and 2400 9.6-second binaural audio mixtures, respectively. Each mixture comprised of two moving speakers and one diotic background noise. The moving speech stimuli were created using the methods described in the **Stimuli Design and Experiment** section. Speech was randomly sampled from the Librispeech dataset^55^. For half of the training data, we chose pairs of trajectories that spanned uniform distribution (quantified by joint entropy); and for another half of the training data, we chose pairs of trajectories whose average distance difference was smaller than 15 degrees to enhance the separation model’s ability to handle closely spaced moving speakers. We randomly chose noise from DEMAND dataset^56^. The SNR, defined as the ratio of the speech mixture in the left channel to the noise, ranged from -2.5 to 15 dB. All sounds were resampled to 16 kHz. We emphasize that the model is speaker-independent as the speakers involved in the testing phase (the voice actors) were not part of the training dataset (Librispeech), ensuring the generalizability and applicability of our system across diverse speakers.

#### Network Architecture and Training

The binaural separation, post-enhancement, and localizer modules were all with a causal configuration of TasNet. For the linear encoder and decoder, we used 96 filters with a 4 ms filter length (equivalent to 64 samples at 16 kHz) and 2 ms hop size. We used 5 repeated stacks witch each having 7 1-D convolutional blocks in the TCN module, resulting in an effective receptive field of approximately 2.5 s. When calculating cos(IPD), sin(IPD), and ILD in Section, we set the STFT window size to 32 ms and the window shift to 2 ms. The binaural separation, post-enhancement, and localizer modules were trained separately. The training batch size was set to 128. Adam^57^ was used as the optimizer with an initial learning rate of 1e−3, which was decayed by 0.98 for every two epochs. We trained each module for 100 epochs.

### Canonical Correlation Analysis

Canonical correlation analysis (CCA) was used to determine the attended talker. From the stimuli side, the inputs involved talker spectrograms and trajectories. We chose a 20-bin mel spectrogram representation obtained with a window duration of 30 ms and a hop size of 10 ms. Audio was downsampled to 16 kHz before mel spectrogram extraction. The mel spectrograms of left and right channels were concatenated along the bin dimension. All trajectories were upsampled to 100 Hz from 10 Hz to match the sampling rate of the neural data. Trajectories were pooled across all trials and normalized. Spectrograms were also normalized on a bin-by-bin basis. We chose a receptive field size of 500 ms for neural data and 200 ms for stimuli spectrograms and trajectories. The starting sample of these receptive fields were aligned in time. Time-lagged matrices were then generated individually for neural data, trajectory and spectrograms.

As done in a previous study^32^, principal component analysis (PCA) was applied individually to time-lagged versions of both spectrogram and trajectory. PCA was also applied to the time-lagged neural data matrix. The top PCA components explaining at least 95% of the variance were retained. This was done to reduce the risk of overfitting in CCA.

CCA filters were trained to project PCA-reduced versions of the attended stimuli and neural data to maximize their correlation. During inference, the trained filters were used to generate correlations for each talker. Attended talker was decided based on voting by the preferences indicated by the first three canonical correlations.

Correction for behavior: For trials in which two or more repeated words were detected in the uncued conversation, the corresponding portions (bounded by button press timings) of the cued to-be-attended and uncued to-be-unattended stimuli were swapped before model training and evaluation. For models trained without correction, no such swapping was done based on behavior.

### Psychoacoustic Experiment

The online psychoacoustic experiment to evaluate system performance was conducted with 24 normal hearing (self-reported) Amazon MTurk participants. These participants were native speakers of American English based in the US. The experiment lasted for a total of 30 minutes per participant and each participant was paid $10. All participants were required to wear stereo earphones.

During the experiment, participants listened to trials and answered questions after each. The task assigned to the participants during the trial was the same as that of the participants from whom neural data was recorded: to attend to/follow the cued conversation (conversation that starts first) and press spacebar upon hearing a repeated word in the conversation being followed. jsPsych was used to design this web-based experiment.

Every participant listened to a total of 15 trials, 5 trials from each of the following conditions: System On (Clean), System On (Separated) and System Off. Neural data was combined from all the three subjects along the channel dimension to test the system. Since a few subjects could have been paying attention to the uncued (to-be-unattended) stream in any given trial, to prevent combining neural signatures across subjects when the subjects were attending to different streams, we discarded trials in which at least one of the subjects had mistakenly attended (tracked at least two or more repeated words) to the uncued (to-be-unattended) stream. This resulted in 18 trials. For every MTurk participant, the selection of 15 trials from these 18 trials and their corresponding condition assignment (SysOn-Clean, SysOn-Sep, Sys Off) were randomized. The order of presentation of these trials were also randomized with the constraint that the first two trials had to be those with Sys Off condition. Throughout the experiment, the participants were unaware of the conditions assigned to the trials.

After every trial, the participants were prompted with the following four questions:

1. Comprehension: A multiple choice question based on the content in the to-be-attended conversation with a single correct answer.
2. Difficulty: Participants were asked to rate how difficult or easy it was for them to follow the cued conversation on a scale from 1 to 5 (1 = very difficult, 2 = difficult, 3 = neutral, 4 = easy, 5 = very easy).
3. Sound Localization: The last three seconds of the trial was allowed to be replayed multiple times by the participants. Participants were asked to indicate from one of five equally partitioned sectors of the frontal half plane (left, front left, center, front right, right) where the cued conversation ended.
4. Voice Quality: Participants were also asked to rate the quality of voices in the cued conversation on a scale from 1 to 5 (1 = bad, 2 = poor, 3 = fair, 4 = good, 5 = excellent).

## ACKNOWLEDGEMENTS

This work was funded by grants from the National Institute of Health’s National Institute on Deafness and Other Communication Disorders (NIH-NIDCD) and the Marie-Josee and Henry R. Kravis Foundation.

## AUTHOR CONTRIBUTIONS

VC, CH and NM designed the experiment. VC and NM analyzed the neural and behavioral data. CH and NM developed the binaural speech separation model. ADM and GMM performed the neurosurgeries. VC, SB and CS recorded neural data. VC, CH and NM wrote the manuscript and made the figures. All others commented and suggested edits to the manuscript.

## DATA AVAILABILITY STATEMENT

The dataset used for training speech separation models and the models themselves are made available with the code on the GitHub repository. The intracranial electroencephalography (iEEG) data that support these findings are also available upon request from the corresponding author (N.M.). These data are shared upon request due to the sensitive nature of human patient data.

## CODE AVAILABILITY STATEMENT

The scripts used to train speech separation models and analyze neural data are made available here: https://github.com/naplab/AAD-MovingSpeakers.

