## Supplementary Document 1 for "Brain-controlled augmented hearing for spatially moving conversations in multi-talker environments"

### SUPPLEMENTARY MATERIAL

| Trial Number | | C1 | C2 | | C3 | C4 | C5 |
|---|---|---|---|---|---|---|---|
| **1** | | Hi Alexander! It is good to meet you. I am Sebastian, and I will be your personal trainer for this month. I am so pumped for our workout today. First, we will lift weights. Then, we will take an aerobics class. For the afternoon, we will go run outside in the park. When we end, I will make a few suggestions for your meals. | Wow. Are you serious? I am still sore from last week's workout. I honestly don't think I have enough energy for an entire day of workout. I already think three days a week at the gym is too much for me. But I also promised myself that I will lose 10 pounds before my birthday this year. I need to change my meals too? Seems like I set the bar too high for myself this year. | | Hi. This is Luigi's Pizza. How can I help you? We have a special two for one offer today. | Hello. I would like to have two pizzas. Can I have one pepperoni pizza and one tuna pizza? Both need to be of medium size. On the second pizza, I would like to have extra olives. I would also like a portion of chips and two cokes, please. Can I have it delivered to my doorstep? The address would be 345 South Street, Apartment A. | For sure! So to sum up, two medium size pizzas, Pepperoni and Tuna with olives, a portion of chips and two cokes? That would be 20 dollars, after the offer. The driver will be there in about 40 minutes. |
| **2** | | Hi Benjamin. You likely know the familiar expression, "An apple a day keeps the doctor away." Although research shows that eating more apples may not actually be associated with fewer visits to the doctor, adding apples to your diet can help improve several aspects of your health. Studies show that eating apples could help reduce stress and slow signs of ageing. What do you think Benjamin? | Hmm. Interesting. Yes, I do agree that apples have great health benefits. Other studies show that eating apples could be associated with a lower risk of several chronic conditions. I eat two apples everyday. I find that eating apples can promote feelings of fullness, decrease calorie intake, and increase weight loss. This might be due to their low fibre content. | | Hello, I have a reservation at this restaurant for two guests at 7:00 PM today. The reservation is on my name, Ms. Lydia. | Welcome to our restaurant, Ms. Lydia. Yes, please follow me. I shall show you to your table. From our records, I see that this is your first visit to our restaurant. As you might know, we specialise in Indian Chinese fusion cuisine. Our appetiser for the day is Manchow soup. Should I get that for both of you? | Yes, please! For the main course, I would like to have a plate of momo dumplings. My partner would like to have a plate of Schezwan noodles. For desserts, could you please get us chocolate ice-cream on honey-fried noodles? |
| **3** | | Hello. I just moved into the city and when I asked around for a gym, I heard a lot of positive reviews about this one. This place is also quite close to my home. I was wondering how much the membership fee at this gym is. Is nutritional advice included in the plan? Could you also mention your opening hours? Also, do you have any personal trainers? | Hi. Welcome to the city. The membership fee is 35 dollars a month. If you opt for an annual subscription, that would be 350 dollars. We always open at 5 AM and close at midnight. We do offer sessions with our personal trainers. For your first classes, we will assign you to one of our expert personal trainers. Nutrition advice is included in the plan. | | Hello, are you Mr. Christopher? Welcome to the UK. I am Alexander and I shall be your chauffeur. Can I help you with your bags? | Hi Alexander. Good to meet you. I am here for a short stay and therefore I don't have any luggages. Do you know a restaurant nearby? I had a long overnight layover at Paris. The restaurants were closed and there was no meal service on the last flight. Oh, another thing! What would be the best time to explore the city? I want to make the most of my short stay here! | Mr. Christopher, the Tom-Tom Sandwich place is a 5 minute drive from this airport. I shall take you there. As the weather is hot, I would recommend that you do sightseeing during evenings. |

| | | | | | | | |
|---|---|---|---|---|---|---|---|
| **4** | | Hi. Are you Elizabeth? Please take a seat. I am Mr. Benjamin Smith, the manager of ABC Software and I would be interviewing you today. First, I would like to congratulate you on passing our aptitude test. You did very well. The recommendations are stellar as well. Could you please tell me how did you hear about this job and what do you know about our company? | Absolutely, Mr. Benjamin. I saw an advertisement online. I clicked on it and it brought me to your company's website where I applied for it. I had read on your website that for over ten years ABC Software has been delivering professional services on software development projects for clients across the globe. I could tell from the client testimonials that you help brands transform their businesses. | | Excuse me, I just arrived in the city today. Do you know where the King's Street is? I am unable to locate it. | Yes, I can tell you where the King's Street is, since I live there. Go straight on and take a left turn at this traffic light. After you turn left, drive straight for half a mile. Make a right turn immediately after you cross the church. That should be the King's Street. I would definitely recommend that you purchase a map to navigate. The city can be quite difficult to drive around for tourists. | Glad that someone could guide me to King's Street finally. Yes, I shall surely purchase a city map. If you are heading to King's Street, feel free to jump in. We can drive you there! |
| **5** | | Hello. I heard that you are the best florist in town. I would like to get my wife some flowers. It is our anniversary today. We have been married for five years. My wife loves roses and daisies, but she really likes any type of flower. I can do traditional, but I think she would like something more creative and unique. Could you please guide me through the best bouquets you have? | Well, happy anniversary! I have some bouquets that combine a few roses with some other seasonal choices. This one here has pink roses with spring tulips and lilies. And it comes with this beautiful vase. We have this vase full of pink and yellow roses. This one here doesn't include roses, but has other gorgeous flowers. It really brightens up the room. | | Good morning! Welcome to our store. We specialise in luxury fashion and apparel. How can I help you today? | Good morning! I was looking for a shirt for my birthday party. Do you have something in blue and medium-size? I liked the one with polka dots that you had put on the mannequin outside. Do you have something as unique as that? I really want to look good at the party. Oh, I almost forgot. I would also like to purchase a pair of shoes that would go with the outfit. | Yes. I think we might have just the right shirt for your special occasion. Could you please try this shirt on? This shirt is from this year's fall collection. The changing room is right over there. If it fits well, I can then take you to the shoes section. |
| **6** | | Hi Elizabeth! When I was a kid, I always dreamt of becoming an astronaut. I used to imagine getting dressed up in a spacesuit, climbing into a spaceship and taking off! But look at how life turned out. I instead became a commercial airline pilot. I still hope that one day I get to fly between planets. Did you also dream of becoming an astronaut one day? | Yes, of course. I did fantasise about becoming an astronaut and floating in space too. But becoming an astronaut is no easy task. One should have also logged in at least a thousand hours of piloting a jet aircraft. But our dreams of floating in space are one step closer, thanks to the new private space companies. Flying between planets would soon become common. | | Hello! How's it going? Sorry, this is my first time ordering coffee at Starbucks. Can you please walk me through the menu? | Sure, I will be happy to show you how to order. For hot coffees, we have Espresso Shots, Cappucinos and Lattes. For cold coffees, we have Iced Mochas, Cold Brews and Iced Espresso Shots. Both the hot and cold beverages come in three sizes: Short, Tall and Venti. You can also customise your beverage with syrups. We can also add foam to the coffee if you need. | Wow. Thank you. That helps a lot. Can I get a hot tall latte with two pumps of hazelnut syrup? A little foam on top would be great. |

| | | | | | | | |
|---|---|---|---|---|---|---|---|
| **7** | | Hi Sebastian, how have you been? Do you know about the new zoo that has opened up in the city? I happened to go there this past weekend. It hosts a unique collection of reptiles, birds and mammals. The most interesting attraction is the dolphin show. It was fun to see the dolphins dance and splash water at the audience. If you haven't been there yet, you must. How was your weekend? | Glad that you enjoyed your weekend at the zoo. I will definitely visit the new zoo. I went shopping on Sunday. My cousins and I are about to go on a trip to Hawaii next month. It has been ages since all of us had hung out together. So we went out to purchase hats and sunglasses. Let me know if you would like to join our trip. We would be happy to have you with us. | | Hi! My name is Alexander. I am a student at Queen's University. I want to rent an apartment and I saw your listing on Oak Street. | Hi Alexander. I am Christopher, the landlord. The apartment was recently renovated. Come inside, let me show you around. It is a one bedroom, one bathroom apartment. This is the bedroom. It is fully furnished with a bed, wardrobe and two bedside tables. There is also a desk for study and a bookshelf. | I love this apartment. It looks cozy and stylish. It is by far the best among all the listings I have visited. Thank you for your time, Mr. Christopher. I shall let you know about my decision to move-in, by the end of the week. |
| **8** | | Hey, I am really hungry. Should we go out to eat? There is a new Italian restaurant that has opened up across the street. They offer pizzas, pasta and spaghetti. My colleagues and I went there last week. We had a good feast. They are also offering discounts for first few opening weeks. After our lunch at the Italian restaurant, we could also go get groceries. | Well, honey, that's a great plan. But let's not go outside today. It is snowing and too cold. Why don't we cook something hot and fresh ourselves? I already brought some groceries on my way home yesterday. I was thinking of chopping the vegetables and making a soup. I can also toast some garlic breads that might go well along with the soup. Sounds delicious, don't you think? | | Hello Ms. Lydia. Thank you for organising this parent-teacher meeting. I am the father of Ethan. | Mr. Sebastian, I could go on and on about how glad I am to have Ethan in my class. He comes prepared to class every single day. When I am teaching, he pays attention and takes detailed notes. He is always at the top of his class when it comes to tests and quizzes. His classmates often ask him for help with classwork and he is always willing to help them. It looks to me like he is doing well both academically and socially. | Wow! Good for him. Well, I appreciate all the positive feedback. I can't wait to go home and tell Ethan how proud I am of his hard work. |
| **9** | | Hey Alexander. I am Natalie. I have a two-bedroom apartment and I am looking for a roommate to share the rent. The address is 378 South Street, Apartment C. It is across from the library and right next door to the pizza restaurant. The rent is 800 dollars per month. The rent is inclusive of electricity, water, internet, and gas. | That seems like a great location. The rent sounds good to me. I also wanted to clarify a few things upfront. I am a non-smoker and I prefer to keep the apartment smoke free. I tend to get up early because I like to get to the gym and work out. My little brother will probably come visit a few times a year. But he is really friendly. I hope these conditions will be fine with you. | | Excuse me, sir. I think you are in my seat. Oh! It's you, Christopher! What a surprise! It's me, Katherine! We were classmates in high school. Do you remember? | Katherine! I am so surprised to meet you here. It has been ages since our graduation. You haven't changed even a bit. You look the same, but better. You didn't show up at our 10-year high school reunion so I haven't heard from you since graduation. Being the smartest person from our class, people were asking for you at the reunion. | Thanks, Christopher. Unfortunately, I wasn't able to make it to the reunion. I am a resource manager at an international travel agency. My work requires me to travel a lot. |
| **10** | | Hey Benjamin. Let's discuss our hobbies. I will start with mine. I have only one hobby. Yet, it consumes all my free time. I enjoy reading and have always been a voracious reader. Although I prefer reading fiction, I make a deliberate effort to read at least one non-fiction | I like to go to the movies a lot. I can watch almost any movie but I am not a big fan of the horror genre. I also like to cook and try new recipes. My mother has always been a wonderful cook, and I feel she is my inspiration. I find cooking very | | Hey Natalie! Thank you for meeting up. I was in town for a short trip and wanted to catch up with you. What have you been up to? | Glad to catch up, Lydia. After high school, I got a degree in finance. I started working at a bank but all along I knew finance was not my thing. I had always been into fashion and clothing, but my parents discouraged me to pursue this career. So I finally made up my mind to go for it. I quit my job | Wow. You rock! I am really happy for you. It takes a lot of guts to start such a business. |

| | | | | | | | |
|---|---|---|---|---|---|---|---|
| | | book in a couple of months. What are some things you like to do? How do you spend your weekends and free time? | therapeutic. On the weekends, I meet friends for lunch, go on hikes, and play video games. | | | and started doing tailoring from home. Then I rented a place and opened a tailor shop. Now I am about to open a boutique with my own creations. | |
| **11** | | Hi Elizabeth. Do you know what would be my dream holiday? A trip to Pairs! Ever since I was a kid, I always dreamt of getting clicked in front of the Eiffel Tower. The city is full of beautiful architecture and museums showcasing fine art from around the world. Having studied French in school, I wish to put it to good use. What would be your dream holiday, Elizabeth? | Paris is definitely a great place. My dream destination would be the Maldives. The resorts and hotels in the Maldives are in a league of their own when it comes to luxury. Every resort has a private beach and allows for recreational activities like scuba diving. You can rent a villa for yourself as well. I have heard that the sea food that gets served in these resorts is delicious too! | | Good morning, students! Please open your books. Alexander, could you please stand up and summarise what we have been looking at, so far? | Sure, teacher. So, we had been studying about the solar system. The solar system consists of the Sun and everything that orbits around it. This includes the eight planets and their moons. The solar system itself is only a small part of a huge system of stars and other objects called the Milky Way galaxy. The Milky Way galaxy is just one of billions of galaxies that in turn make up the universe. | Thanks, Alexander. That was a great summary. Today, we shall take a look at our home planet, Earth. Earth is the third planet from the Sun. |
| **12** | | Elizabeth, I was just thinking about my pet, Rosco. He is from the breed of German Shepherds. He is 2 years old, fluffy and black in colour. When we return, he gets delighted to see us and wags his tail rapidly. The only inconvenience we have is that he snores very loudly. So we put him to sleep in a different room. Lydia, do you have a pet? | Yes, Katherine. I am more of a cat lover. I have a pet cat named Ginger. She is white in colour, has beautiful eyes and a fluffy hair coat. My uncle gifted Ginger to me on my twentieth birthday. Sometimes, she can be lazy. She is quite friendly with dogs. We should definitely let Rosco and Ginger meet each other. They can become good friends. | | Hello. I need to get to New York City by 5 PM today. Is there a train that can get me there by that time? | Yes. We have a train leaving in 30 minutes that will get you there at 2:45 PM. Or, if you prefer to leave later, we have an express train that leaves at 3:00 PM and arrives at 4:55 PM. You can purchase a ticket either for the first class or the second class. The first class ticket costs 100 dollars, but it includes a meal and a drink. The second class costs 50 dollars. | Thanks. I will take the earlier train with a second class ticket. I would rather leave early and be relaxed, than leave later and be worried that I might be late. |
| **13** | | Lydia, have I told you about my hometown? I come from a small village situated in Austria. It is a very quiet place, the air is clean and fresh. The landscapes are breathtaking. What I like the most about my hometown is that you can enjoy several activities, such as mountain biking and hiking. I go hiking very often with my friends. Lydia, tell me something about your hometown. | For sure. I come from New York City. I live in a three-bedroom apartment on the eighth floor. My school is not very close to where I live, so I have to take the subway. You can reach almost every part of the city easily with the subway. I like New York because there is so much great food, everywhere. So many places and cuisines to try. | | Hi Sebastian. You are very late. What happened? I was so surprised when you called me yesterday and told me you wanted to come to the gym with me. | I am so sorry. I would have been on time if the traffic hadn't been so bad. You know, lately I have felt so out of shape, so I want to change my lifestyle. And I want to lose weight, too. If I had not gained so much weight, I would have been more energetic. I know you are an expert in this. So if you have any advice for me, I would be happy to listen to it. | Well, if you didn't eat so many sweets anymore, you would definitely lose weight. You should also try to avoid junk food. Eat lots of fruits and vegetables. |

| | | | | | | | |
|---|---|---|---|---|---|---|---|
| **14** | | Hey Natalie. Some people say academic excellence is the only requirement for a successful career. To me, this seems to be a very narrow-minded view. Degrees alone cannot guarantee success. They may help land you a job, but, your success depends upon how you think, behave, talk and present yourself. It is one's attitude towards life that brings success. | Yes, Alexander. You are absolutely right. While accumulation of knowledge is definitely good, what's even more important is its application. Successful people are those who are passionate and use their knowledge to create value in the world. For example, a self-taught programmer, without formal schooling, can write an app that would top the charts and make him millions of dollars. | | Wow! This burger is so good. I think the food here must be different from what you eat in Spain. Are you still enjoying living in the US? | Yes! Sometimes I get a little homesick, though. I also get tired of speaking English all the time. But actually, I am really proud of the way I have handled the cultural differences. Just the other day, I was getting ready for class. I was amazed by how comfortable I feel with a stricter schedule now. In Spain, we like to take our time. Here, in the US, you need to be on time everywhere you go. | I am glad that you have grown accustomed to your new schedule. I am excited about going to our school's football game today. Would you like to join? |
| **15** | | Christopher, this is the third time you have been late this week. You were needed in the meeting today. Fortunately, the deal was done, but it almost did not happen. If you keep getting stuck in traffic everyday, you should probably plan to leave your house early. Hope you won't be late to another meeting. You must be on time as a working professional. This is not acceptable. | I am sorry. Today, it wasn't the normal traffic rush. I was driving to work when my car was stopped by a policeman. I was pulled over, and the policeman told me I was under arrest. He said that my car was at the scene of a crime. Then, the policeman got a call on his radio that the actual robbers had been caught. After hearing this, the policeman let me go. | | Hey, Sebastian! It must have been a year since you began working in the United States. Is there anything you like better in the US than in France? | Well, I have had loads of fun taking long road trips. There are some beautiful places in the US that I didn't know about. One of my friends lives in Boston. She took me home with her when we had a study break. I was fascinated by the beautiful mountains we drove through. I was a little nervous about meeting her family, though. But it turned out that her mother is married to a French man! | Wow! What a coincidence. Living in another country must be challenging. I am really impressed by how you have managed your stay here so far. |
| **16** | | Alexander, I think the most pressing question of the decade is whether robots will replace humans. We have all grown up watching movies such as The Terminator, Star Wars amongst many others. Such movies have rooted in us the fear of being replaced by robots. This fear of being the subservient one has considerably increased the research into this field. What do you think? | Yes, I agree. Every day there is a new article on technology changing the way we work. Such articles are usually accompanied by a list of occupations that will be replaced by robots, and the skills that will be irreplaceable in the world dominated by machines. I don't think a robot will be able to mimic human emotions and offer empathy in complex environments. | | Hi Elizabeth. It is good to see you finally. Every time I suggest a meeting you always have some excuse. Are you trying to avoid me? | No, no! You know I always look forward to meeting you. I have been meaning to pay you a visit, but I just couldn't find the time. I have been very busy with my job. I want to get promoted and earn more, but promotions take a lot of work. Imagine having to get up at 5 AM everyday and going to sleep at midnight! I aspire to become the CEO in the next five years and I am working towards this. | If you continue to work so hard, you risk getting sick. You also never get to live life to the fullest and spend enough time with the people you love. |

| | | | | | | | |
|---|---|---|---|---|---|---|---|
| **17** | | Hi. I need to get to the airport. Do you have a direct metro line to the airport? I am late for my flight and I need the quickest possible option. I tried to book a taxi but none seem to be available today. I would have taken the bus, but buses are quite slow, with many stops in between. So I desperately hope that there would be a direct line to the airport. | Sorry, we do not have a direct metro line to the airport. But you can easily get there by making just one transfer. Take line 10 to South Street. Then, you should change to line 8 and ride it all the way to Dayton International Airport. It takes about 10 minutes to get to South Street. The line 8 ride takes about forty minutes. This should be the fastest option currently. | | Hey Elizabeth. Have I told you that I want to apply to the Academy of Dramatic Arts and become an actress? | Wow. That sounds really awesome. Well, if I had good looks and any acting skills, I would have considered becoming an actress too. The only problem is that my dad is not supportive. He said that if I became an actress, he would be extremely disappointed. So, I now work at a bank. I like my current job, but I believe there would have been more grounds for expression if I had been an actress. What would you do if you were in my place? | Oh, I am very sorry to hear that. If I were you, I would try to change his mind. I would express all my feelings and make sure that he really understands. |
| **18** | | Natalie, what is your favourite genre of music? It is hard to pick a favourite kind of music for me because I love all music. I usually pick my music based on the mood. When I am stressed, I listen to classical music. The instruments that play in a classical piece create a peaceful melody that helps me relax. Sometimes it gets me relaxed to the point that I fall asleep. | Hmm. That's actually funny. My favourite genre is electrical dance music. The reason why I like it is because I like beats that are catchy but not too catchy. This genre is best suited for parties. It gets everyone at the party in the mood for dance. I also like jazz. I only listen to it whenever I need to concentrate on a particular task. | | Hi Benjamin! It is so good to see you. How are you doing, son? Have you settled in? How do you find your new apartment? | Hi Dad. I am doing good. The new apartment is great and I love it. It is cozy and convenient for getting to college. And my roommate is very nice as well! My bedroom is very large and has a small closet. I keep it very tidy. The kitchen is old, but all the appliances still work fine. The only downside to this apartment is that I have to share my bathroom with my roommate. | Wow. Your apartment looks nice. I hope that you will get along well with your new friends. Your mom and I shall surely pay a visit some day. Talk to you later. Bye. |
| **19** | | Hi there. I was wondering if you could help me. I am not from the city and I am trying to use the bus system for the first time. My plan for today is to visit the modern art museum, stop by the zoo, go to the rose garden and then eat dinner downtown. Is there a bus pass that I can purchase? And which bus should I catch for the museum? The map on the board is too confusing. | Welcome to the great city of Chicago! I can certainly help you navigate the bus routes. You should probably get a 24 hour bus pass for unlimited bus rides. That way, you can stop and explore as much as you want in each neighbourhood for the entire day. Just show it to the driver each time you board the bus. To head to the museum, catch the number five bus. Enjoy your day! | | I am so excited to visit New York City next week. I will be staying there for over a month. | You are so lucky! I always wanted to go there. I wish I could go with you. But I will be babysitting my cousin's children next week. My cousin has an important meeting that he needs to attend. I believe New York City is the most linguistically diverse city in the world. The last time I checked, there were more than 800 languages being spoken in NYC. It is also home to more than 8 million people. So, what will you be doing there? | I will be seeing all the sights. I will be touring everything from the Empire State Building to the Statue of Liberty. I am sorry that you cannot come but I will send you pictures. |
| **20** | | Hey Lydia! Last summer, I visited my grandparents' place in the countryside with my parents for a couple of weeks. Mostly, we went hiking in the hills and mountains nearby. We also just hung out in the village, playing cards and eating. Our grandfather loves | My family and I went to Hawaii last summer. The trip lasted 10 days. Our first stop was at Maui, where we went whale watching. Our second stop was at Pearl Harbour in Honolulu. The final stop was at Mauna Loa, one of the world's most | | Hi! Are you a new exchange student? I saw you in my Math class. How has your first month in the US been? Hope you were able to explore the city. | Yes. I am a new exchange student from Spain. I am studying at Boston University for a year. I am having a great time. I am glad that I made the decision to study abroad. I saw Boston in some American movies. It looked so nice that I wanted to make the | I hope you get to take a break soon. There are so many things to see in Boston. I might go out this weekend to do some shopping for my apartment. Let me know if you want to tag along. |

| | | | | | | | |
|---|---|---|---|---|---|---|---|
| | | gardening and we helped him plant saplings and water the plants in the garden. Lydia, Where did you go last summer? | active volcanoes. We plan to go back to Hawaii this summer. Let me know if you would like to join us. | | | effort to come here. I wanted to go sightseeing, but I have been too busy doing my homework. | |
| **21** | | Benjamin, you play a lot of video games, don't you? Yesterday, I read an article discussing if video games should be considered a sport. Aside from improving hand-eye coordination and fine motor skills, the article claimed that video games also teach puzzle-solving skills and can be a valuable tool for helping children to learn teamwork. Benjamin, do you agree with this view? | I agree with it to some extent. I play a lot of video games. Studies even show that regularly playing video games improves spatial recognition, multitasking skills, problem-solving skills, and perception. Since video games lack the typical physical elements found in sports such as Rugby and American Football, many argue video games aren't the same thing. | | Wow. Can you just believe our high school is over? I feel like time just flew by. Natalie, where do you see yourself ten years from now? | I believe I will be a very famous basketball player. By the time I finish college, I will have mastered all of the skills I need to succeed as a pro player. I will have worked with great coaches and played against tough teams. I will have joined an incredible team by then and won championships year after year. I will have turned into an amazing athlete. Just wait. You will see. | I believe in you, Natalie. If anyone can do it, you certainly can too. Anything is possible. Whatever the next ten years hold, we know that it is going to be exciting. |
| **22** | | Ms. Elizabeth, thank you for coming to today's parent-teacher meeting. I really wanted to speak with you about George's behaviour in class and grades. His classroom behaviour has really been a problem. He doesn't pay attention when I am teaching and he often tries to distract his classmates. I ask him to stop distracting and this usually gets him to focus for a few minutes. | I am extremely sorry to hear that. I honestly wish he would just behave well in the first place. I was thinking that he might respond well to a reward, like 15 extra minutes of recess for good behaviour. I think he can really improve if he puts in a little more effort and focuses more in class. I will talk to him tonight. Ms. Katherine, thanks for bringing this to my notice. | | Hey, Alexander! I have been wondering how often all of us exercise these days. I go running thrice a week. How often do you exercise? | I hardly ever go to the gym, but I have an active lifestyle. I usually cycle to work. Sometimes I take the stairs instead of the elevator. If I have groceries, instead of taking all the bags in one round, I take two bags at a time, and go up and down several times. In the evening, I normally do the dishes, dust, and mop. I use housework as a substitute for sport. Cleaning is great body work for me. | That's nice. You definitely seem to have a healthy lifestyle. I hope that you maintain your lifestyle that way. |
| **23** | | Hi Alexander! Have you ever wondered how can one always stay happy? Regardless of one's version of happiness, living a happier life is within reach. A few tweaks to regular habits can help get there. I find exercise and dancing to be instant happiness elevators and great stress relievers. Do you have any techniques that you use to keep yourself happy? | Yes indeed. While life can be a jittery roller-coaster ride, I do believe a few techniques can help achieve peace and happiness during rough times. A simple technique that has helped me a lot is expressing gratitude! Simply being grateful to things and people can give your mood a big boost, among other benefits. I start each day by acknowledging one thing I am grateful for. | | Hi there! I am Christopher and I am a passenger travelling on the flight CA 187 to Los Angeles. Is this the counter for Catalina Airways by any chance? | Yes! This is indeed the right counter for your flight. We were just about to close our counter. I see your information here. If you have any baggages, could you please place them on the weighing scale? I see that you have chosen a window seat. Let me print out your boarding pass and you will be ready to go! Okay, here it is. You are going to depart from Gate 54 at Terminal C. Enjoy your flight! | Thank you so much! This is actually my first flight and I am very excited about it. I shall now head to security. |

| | | | | | | | |
|---|---|---|---|---|---|---|---|
| **24** | | Elizabeth, yesterday I read an article discussing the pros and cons of social media. I believe the pros outweigh the cons. Social media has removed all barriers to communication by bringing the globe on a single platform. From emergency alerts and announcements to knowing how our friends are doing, gathering information has become so convenient. Don't you think so? | Unfortunately, I do not agree completely. It takes a tremendous amount of effort and discipline to stay away from social media. Results have shown that it has had adverse effects on human minds and their functioning. Outdoor activities among children have also reduced significantly. It is hard to ignore the vast amount of fake news that spreads like wildfire through social media. | | Natalie, your dad and I find you to be very upset these days. Is someone or something bothering you at school? | Mom, my friend got a new smartphone. Her smartphone is the slimmest smartphone on the market. It is much slimmer than mine. My phone is older than all the other phones I see. It is also super slow. It takes more than two minutes to load an app. I have been using the same phone for the past four years. I wish I had a more modern phone. | Natalie, a phone is the least important thing in life to make you happy. A more modern phone won't make you happier. But, yes. I really think that you need an upgrade. Why don't we go to the store today and buy you a new phone? |
| **25** | | Hey, Sebastian! Yesterday, I was reading about an article on global warming. It got me thinking about the way we lead our lives and how it impacts our planet. I think it is high time that we start thinking about becoming carbon neutral. I have already installed solar cells on my roof. Sebastian, what do you think are some steps one can take to reduce his carbon footprint? | Well, I am sure there are plenty ways. One easy way to cut down on carbon emissions would be to replace the usual tube lights in our homes with LED bulbs. LED bulbs use way less energy and are cheaper. Using paper bags instead of plastic bags, planting more trees and driving fuel-efficient vehicles can also help cut down a person's carbon footprint. | | Excuse me, I have a reservation at this hotel for a stay lasting 4 nights and 5 days. Would it be possible to get a room on the 9th floor with a single bed? | Hi Mr. Alexander. Your company had informed us about your stay. Yes, I can give you a room on the 9th floor with a single bed. Here is your key. It is room 925 facing the sea. Oh! One other thing. The stay package comes with complimentary breakfast for all mornings. If there are any concerns with your stay, please dial 1 from the room telephone. | Thank you so much! I read a lot of positive reviews about your hotel on the internet and I am indeed excited for my stay here. I shall let you know if there are any concerns. |
| **26** | | Katherine, I am curious to know how our daily schedules look like. I get up at 7 AM and have breakfast with my family by 8 AM. We all leave home by 9 AM. My parents drive me and my little sister to school and then they go to work. I take lunch at the cafeteria at noon. I finally reach home at 6 PM. Are your days as busy as mine too? | My schedule is more or less the same. My day starts at 8 AM. I have breakfast just with my dad. Then, I go to school by bus. After I finish school, I go straight to my home. I do my homework at 3 PM. At 5 PM, I go to a dance club where I do yoga and take dance classes. Before going to sleep, I like reading. | | Hey Lydia. I just realised that I have never met your family. Do you have a big family? | Yes, I indeed have a big family. I have two twin brothers, a sister, a niece, a brother-in-law, and there's my mother and my father. I take after my dad in appearance, and after my mom in personality. My mother is short and plump, has brown hair and big blue eyes. She is really beautiful. My dad is as tall as me, has grey hair, brown eyes and a moustache. He is pretty quiet and doesn't talk much. Would you like to meet them next weekend? | I would love to! Thanks for inviting me. You are so lucky to have such a wonderful family, Lydia. |
| **27** | | Benjamin, I hope your classes at university are going well. It is nice to talk to you on a video call. I can see that you have settled in your new apartment. I also heard that it is a great location. Tell us about the part of the city you live in. Are there any | Oh, dad! You are always thinking about food! Actually, yes! My roommate and I have eaten at many restaurants nearby. There is a really great Indian restaurant. It's a small restaurant but they make really good food. Then, there is | | Please get in. Where are you headed today? Are you in town for business or is this a vacation? | Thanks. I am headed to the Hilton Hotel. The one on 21st street. It is both a vacation as well as a business trip. I have a meeting to attend for work. But I also have time to relax and enjoy the city for a few days. This is my second time in this city. | My favourite is Nathan's Diner on Elm Street. They have the best cheeseburgers. Here we are. The Hilton Hotel. This is the newest hotel in the city. I have heard it is a great place to stay. |

| | | | | | | | |
|---|---|---|---|---|---|---|---|
| | | good restaurants to hangout and eat? Is there anything fun to do nearby your apartment? Have you and your roommate gone out recently? | this venue nearby where different musicians perform. We have seen some cool bands play there. They use fog machines and really loud speakers. | | | I visited here with my family a long time ago. A lot has changed since then. Any recommendations for restaurants I should try? | |
| **28** | | Natalie, I am sure we all have people in our lives who have played an influential role. In my case, the most influential person in my life is my father. He is always pushing me to be the best. Whether it is with school, sports, or even a silly competition, he wants me to be the best. Natalie, who has been the most influential person for you? | That's awesome, Lydia. Glad to hear that. In my case, my mother has been an extraordinary influence on my life. The one trait of hers that I value the most is her enthusiasm for learning. There has never been a night I went to sleep without learning something new from my mother. It is this never ending passion to learn that has made me curious and question things around me. | | Do you like living here, Elizabeth? If not, what do you think would be your ideal home? | Are you kidding? I hate this place. I am always surrounded by traffic and other annoying noises. Not to mention there's so much dust and pollution here. I hope to be able to move out of the apartment soon. My top choice would be a big house in the countryside, with a view of the village, trees and fields. It has always been my dream to live in a quiet, peaceful place. What would be your ideal home, Katherine? | Well, I am more of a city girl. My ideal home would be an apartment in a big city like New York. Right downtown in the heart of it all. Things would be happening all the time and I wouldn't be bored. |

Table S1: Transcripts for the conversations used in the trials. C1 and C2 correspond to the utterances of the talkers in the cued conversation. C3, C4 and C5 correspond to talkers in the uncued conversation.

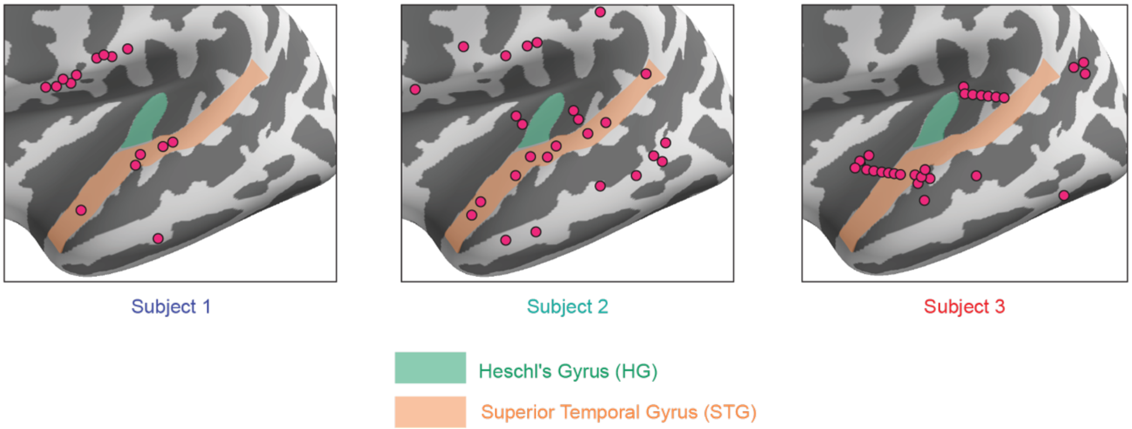


Figure S1: Sites of speech-responsive electrodes used for analysis from the three subjects. All subjects had coverage over their left temporal lobe.

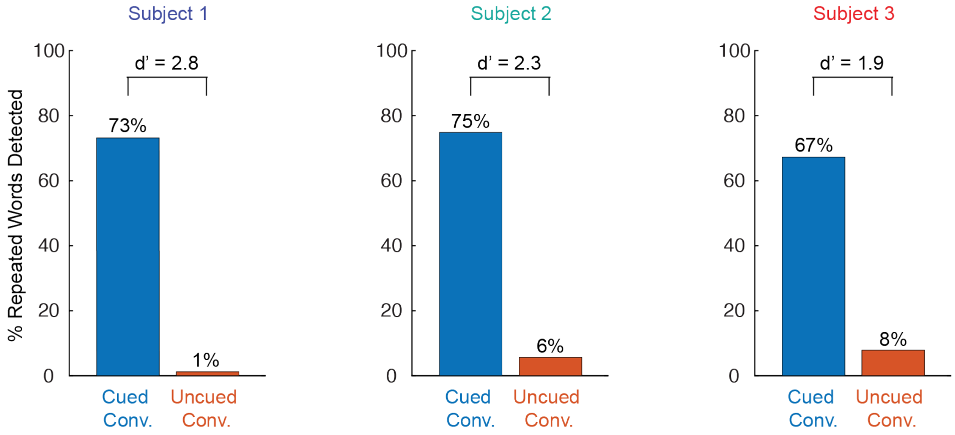


Figure S2: Proportion of repeated words detected in the cued (to-be-attended) and uncued (to-be-unattended) conversations by the subjects across all trials. Subjects mostly attend to the cued conversation, but sometimes they also pay attention to the uncued one.

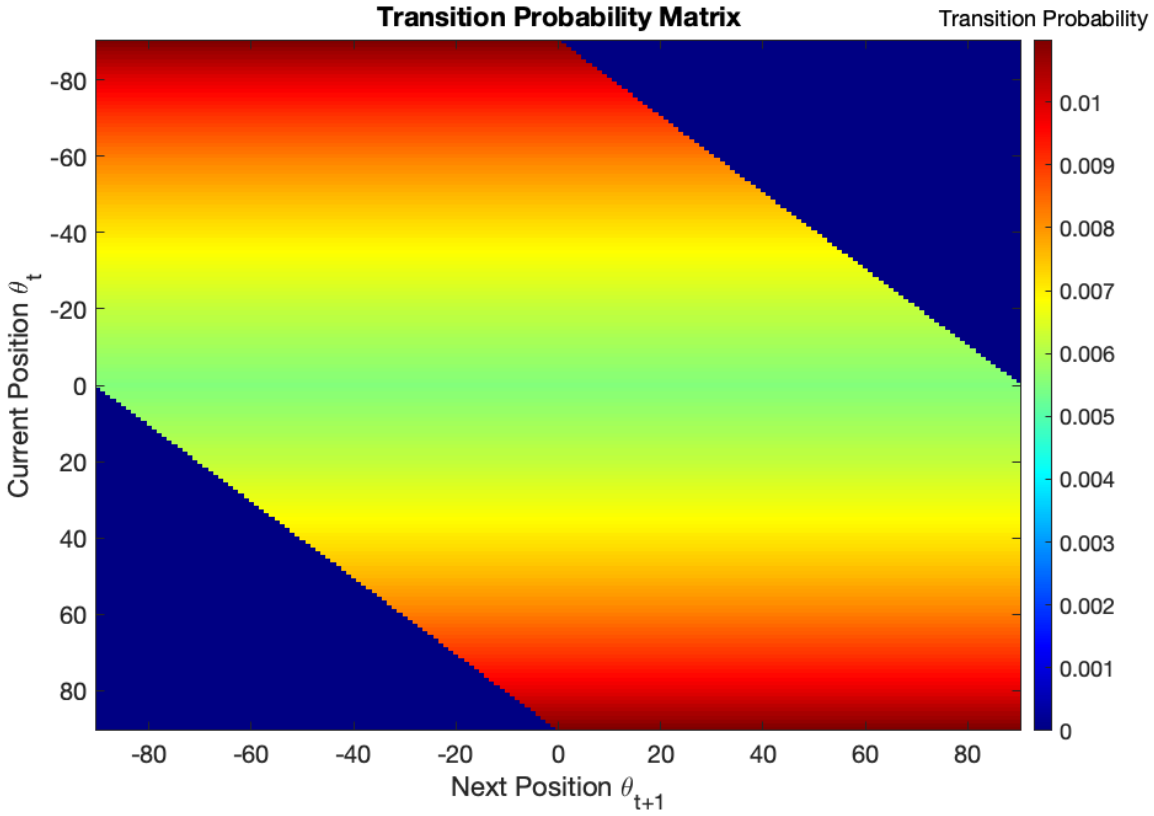


Figure S3: Transition probability matrix determining the trajectory generation process using a first order Markov chain.
